# High resolution 3D imaging of liver reveals a central role for subcellular architectural organization in metabolism

**DOI:** 10.1101/2020.11.18.387803

**Authors:** Güneş Parlakgül, Ana Paula Arruda, Erika Cagampan, Song Pang, Ekin Güney, Yankun Lee, Harald F. Hess, C. Shan Xu, Gökhan S Hotamışlıgil

## Abstract

Cells display complex intracellular organization through compartmentalization of metabolic processes into organelles, yet neither the resolution of these structures in the native tissue context nor its functional consequences are well understood. Here, we resolved the 3-dimensional organelle structural organization in large (>2.8×10^5^μm^3^) volumes of intact liver tissue (15 partial or full hepatocytes per condition) in high resolution (8nm isotropic pixel size) by utilizing enhanced Focused Ion Beam Scanning Electron Microscopy (FIB-SEM) imaging, followed by deep-learning-based image segmentation and 3D reconstruction. We also performed a comparative analysis of subcellular structures in liver tissue of lean and obese animals and found marked alterations particularly in hepatic endoplasmic reticulum (ER), which undergoes massive structural re-organization in obesity characterized by marked disorganization of stacks of ER sheets and predominance of ER tubules. Finally, we demonstrated the functional importance of these structural changes upon experimental recovery of the subcellular organization and its marked impact on cellular and systemic metabolism. We conclude that hepatic subcellular organization and ER’s architecture is highly dynamic, integrated with the metabolic state, and critical for adaptive homeostasis and tissue health.

---

Organs and tissues exhibit distinct features of structural organization at various scales to meet functional demands and adapt to challenges to maintain homeostasis and viability^1^. At the subcellular level, metabolically specialized cells also present distinct morphological and spatial organization of organelles^2,3^, which likely plays a role in compartmentalizing and partitioning of metabolic processes, to increase functional diversity and provide metabolic flexibility. Therefore, the intracellular architecture of organelles is directly related to the cell’s specialized function and it’s critical to explore intracellular structural properties in their native tissue environment to understand structure-function relationship in health and disease. A major challenge is the extreme difficulty in resolving organelle structure and organization in the native tissue context in vivo, especially for organelles with complex architecture such as endoplasmic reticulum (ER). This is in part due to technical limitation to study at high spatial resolution in large volumes of tissue with the existing and commonly used microscopy approaches and analytical platforms. Most of the studies on organelle architecture and cellular organization have been performed in model organisms such as yeast or cultured cells and provided valuable information^4–8^. However, specialized cells of mammals in their native tissue environment, present a more complex organization that may not be fully and accurately captured in these systems. For example, cells that secrete high amounts of proteins such as pancreatic acinar or plasma cells are filled with rough ER in the form of stacked sheets^2^, whereas steroid/lipid hormone secreting Leydig cells have a vast network of smooth tubular ER^3^. Cells in these classes, generally execute one predominant function such as protein or lipid synthesis and exhibit a more, static or stable homogenous structural state of the ER to support that particular function. On the other hand, cell populations such as hepatocytes are charged with pleiotropic metabolic functions and present highly heterogenous structures to, presumably, support their multifunctionality.

In order to establish structure-function relationship between organelle architecture and metabolic function in homeostasis and disease, we chose to study hepatocytes in their native liver tissue environment. To capture the full resolution of in vivo hepatocyte subcellular structures in liver and explore a comparative analysis of normal and obese tissue, we took advantage of enhanced FIB-SEM imaging, a technique that uses scanning electron microscopy to scan the surface of a volume, followed by milling of the surface with a focused ion beam. Recent advances in FIB-SEM imaging technology^9–11^ substantially expanded the imaging volume while maintaining isotropic nanometer resolution, thus allowing us to image large intact liver volumes derived from lean and obese animals at voxel size of 8 nm in x, y, and z dimensions.

Using this approach, we obtained a volume of 96μm(x), 64μm(y), 45μm(z) comprising 5638 consecutive images for the liver derived from lean mice, (**Fig. 1a, c, Supplementary Video 1**). For the obese condition, we obtained liver volumes from two different obese mice. The first sample consisted of 81μm(x), 73μm(y), 63μm(z) comprising 7896 consecutive images (**Fig. 1b, d, Supplementary Video 2**) and the second sample was 64μm(x), 64μm(y), 68μm(z) comprising 8501 consecutive images (**Extended Data Fig. 1a, b**).

**Fig. 1.**
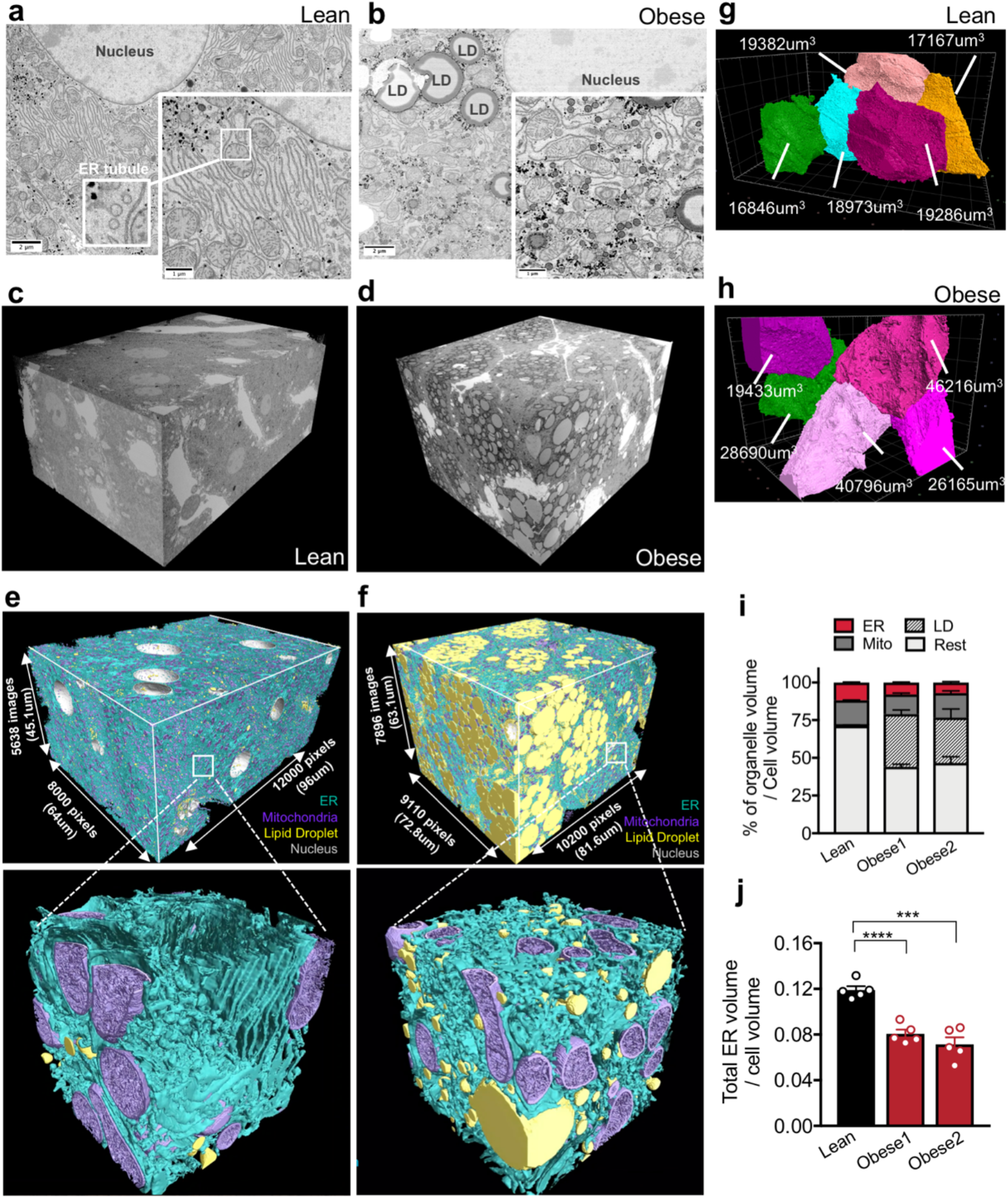
FIB-SEM imaging and automated deep-learning-based segmentation of organelles from intact liver tissue derived from lean and obese mice. **a, b,** Single section SEM of liver from lean and obese mice liver in fed state at 8 nm pixel size. Inset, in **a,** highlights the tubular ER structure. **c, d,** 3D reconstruction of FIB-SEM images derived from liver volumes from lean (**c**) and obese mice (**d**). (**Supplementary Videos 1, 2**). **e, f,** Convolutional neural network based automated segmentation of liver volumes derived from lean (**e**) and obese (**f**) mice. The dimensions of the volumes are depicted in the figure. ER (endoplasmic reticulum, blue), Mito (mitochondria, purple) LD (lipid droplet, yellow), Nucleus (gray). Inset images show 500×500×500 pixel^3^ magnified volume from the whole datasets (**Supplementary Videos 3, 4**). **g, h,** Reconstruction of 5 full or partial hepatocyte volumes present in the liver volume imaged by FIB-SEM. The volumes of the cells are depicted in the figure. All the reconstructions were performed in Arivis Vision 4D software. **i**, Quantification of total ER volume normalized by cell volume in 5 different cells derived from lean and obese mouse 1 and obese mouse 2 (***p=0.0001). **j**, Percent of organelle volume normalized by total cell volume; n=5 cells for lean and for 2 obese.

An additional major challenge using this method is the identification, segmentation and quantification of each sub-cellular structure within the cell they belong as it is not possible to achieve this task in a high-throughput manner with manual annotation or the standard segmentation techniques such as thresholding, K-means clustering or edge detection. In order to overcome this challenge, we took advantage of machine learning based approaches and utilized convolutional neural networks to automatically segment organelles (ER, mitochondria, cristae, lipid droplets, peroxisomes and Golgi), plasma membrane and the nucleus in a volume of 540 billion voxels in lean and 730 billion voxels in obese liver (**Fig. 1e, f, Supplementary Videos 3, 4, Supplementary Table 1**), following the pipeline in **Extended Data Fig. 2a, b**. Cloud computing has been utilized to handle the computation needs of the terabyte-size datasets and the success rate of segmentation for all classes was measured by the Jaccard Index or evaluating the object-level precision and f1-score. In order to exclude the non-cell volume occupied by blood vessels and sinusoids, we quantified morphological characteristics of the segmented organelles in the 5 largest cell volumes in each dataset as shown in **Fig. 1g, h, Extended Data Fig. 1d.** A whole hepatocyte volume from the lean tissue was around 19×10^3^μm^3^, whereas obese cell was 40×10^3^μm^3^. Using this approach, we observed that the lipid droplets content was less than 1% in the cells from lean mice and increased dramatically to 30 to 34% in the obese cells. The mitochondria volume corresponded to 16% in lean and 13%-16% in obese cells (**Fig. 1i**). The total ER volume corresponds to ~12% of total cell volume in the lean hepatocytes and decreased to ~5%-8% in the hepatocytes from obese mice (**Fig. 1i**). However, when we normalized the total ER volume present in the cells per cytosolic volume only, we verified that lean and obese cells present comparable ER volumes (**Extended Data Fig. 2c**). These results indicate that obesity does not alter the amount of total ER volume per se and the apparent decreased volume ratio can be attributed by the increase in cell size due to lipid droplet accumulation.

Although the apparent volume of ER was comparable between lean and obese samples, we noticed striking differences in the organization and composition of the ER membranes in the cells. This led us to focus our analysis to the ER in greater detail. Morphologically, ER comprises a continuous network of interconnected membranes subdivided in the perinuclear ER, which wraps the nucleus and the peripheral ER, which can be seen as a network of tubular shaped membranes devoid of ribosomes (smooth ER) along with stacked flat sheets formed by two flat membrane bilayer characterized by the presence of ribosomes (rough ER)^12–15^. As shown in **Fig. 2a, b, Supplementary Videos 5-6,** through our imaging and reconstruction, we obtained an unprecedented resolution of ER sheets and tubules which have not been possible to be resolved in native large tissue setting with traditional imaging methods available before. Using this approach, we observed predominance of the well-organized parallel sheets of the ER in cells from lean tissue and the dramatic reduction and disorganization in these structures in obese liver, along with the enrichment in tubules (**Fig. 2c, d, Supplementary Videos 7-9**). In order to quantify the transitions of ER subdomains in these conditions, we re-segmented the ER volume separating the ER sheets from tubules. For that, we manually labeled around 20 consecutive binary ER mask images, generated grand truth and utilized a U-Net neural-network-based approach to separate ER into these 2 sub-domains (**Fig. 2e, f**). As shown in **Fig. 2g and Extended Data Fig. 2d**, the percent of ER sheets per total cell and cytosolic volume is significantly decreased in obese hepatocytes while the ER tubules are significantly increased per cytosolic volume (**Extended Data Fig. 2e**). In liver tissue derived from the lean mice in the fed state, the ER sheets corresponded to 30-35% of total ER volume (**Fig. 2h**). The ER sheet/tubule volumetric ratio corresponded to 0.49-0.55 (**Fig. 2i**) and surface area ratio corresponded to 0.5 (**Extended Data Fig. 2g**), reflecting these cells’ synthetic and anabolic state. In obesity however, we detected a significant reduction of ER sheets that correspond to ~18 to 20% of the total ER volume (**Fig. 2h**) and a markedly decreased sheet/tubule volumetric ratio of 0.25 (**Fig. 2i**).

**Fig. 2.**
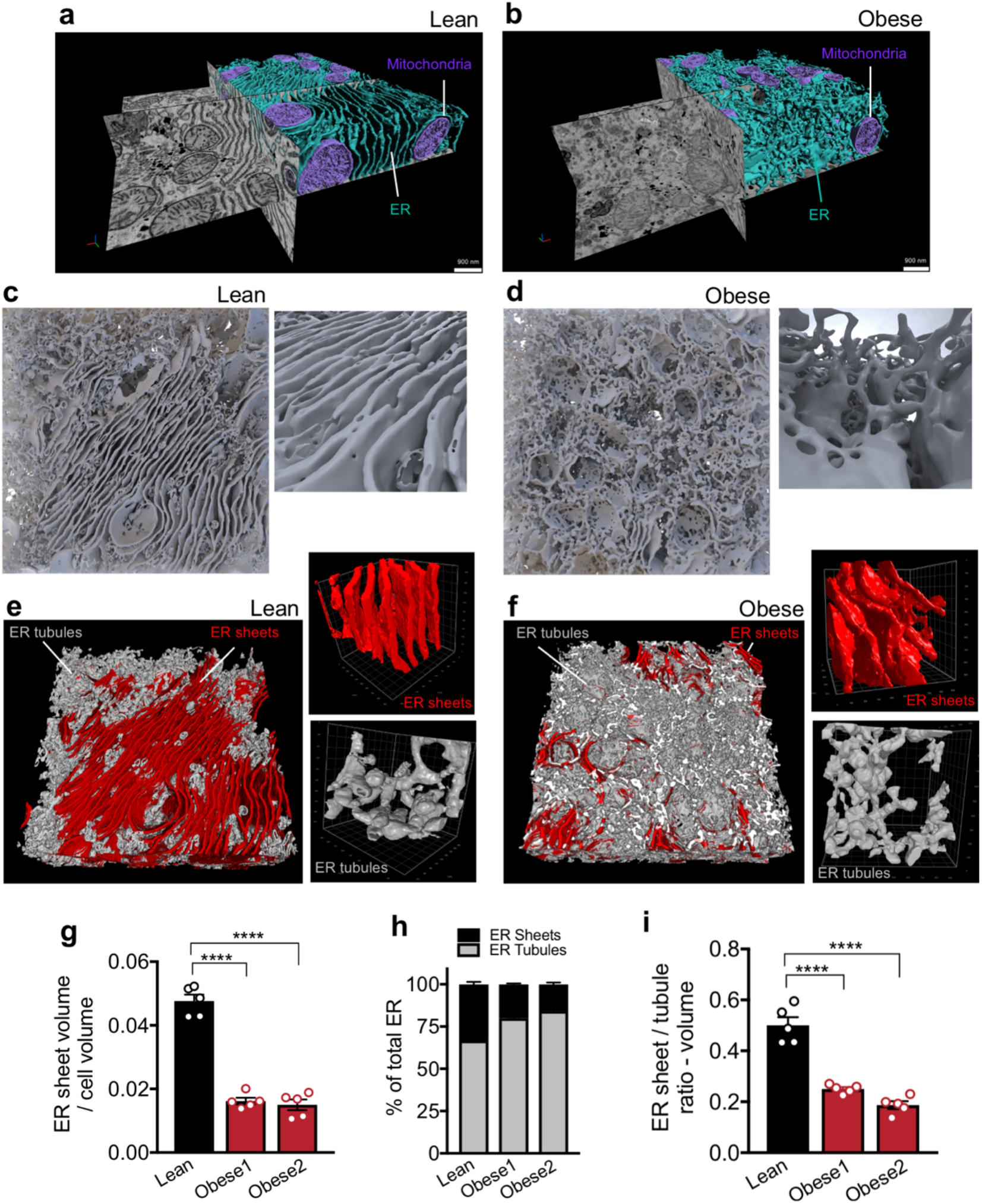
Large tissue FIB-SEM imaging reveals hepatic ER sheet / tubule ratio is decreased in livers of obese mice. **a, b,** Partial reconstruction of segmented ER and mitochondria on raw FIB-SEM data derived from hepatocytes from lean (**a**) and obese mice (**b**) (**Supplementary Videos 5, 6**). **c, d,** 3D reconstruction of segmented ER morphology from lean (**c**) and obese (**d**) liver (1000×1000×400 pixels – 8×8×3.2um^3^). 3D reconstruction images were generated using Houdini (SideFX) software. Inset shows the ER sheets and tubules in higher magnification (**Supplementary Videos 7-9**). **e, f,** Sub-segmentation and 3D reconstruction of ER sheets (red) and tubules (gray) from lean (**e**) and obese (**f**) mice. Magnifications show 100×100×100 pixel^3^ representation of ER sheets and tubules separately. **g,** Quantification of ER sheet volume normalized by cell volume. n=5 cells for the 3 datasets (****p<0.0001). **h,** Quantification of percent of ER sheets (red) and tubules (black) relative to total ER from lean and obese cells. n=5 cells in each group. **i,** Ratio between ER sheets and tubule volume (****p<0.0001).

In addition to the decreased abundance of ER sheets, we also observed that the organization and parallelism of the sheets were disturbed in obesity (**Fig. 2e, f insets**). In order to quantify the parallel organization of ER sheets in higher number of biological replicates, we performed TEM images from livers, where the whole hepatocyte mid-cross-sectional area was captured (around 15 full hepatocytes per condition). We manually annotated the ER sheets, generated binary ER masks and applied an algorithm to segment the ER traces that are found in parallel-arrayed organization (**Extended Data Fig. 3a).** We validated the algorithm using acinar cells as a positive control, given that these cells are strongly enriched in parallel organized ER sheets **(Extended Data Fig. 3b)** and a mouse hepatocyte cell line (Hepa1-6) as a negative control, since these cells are devoid of parallel organized ER sheets **(Extended Data Fig. 3c)**. Based on this algorithm, we validated that obesity leads to a reduction in parallel-stacked organization of ER sheets in regard to the total ER (**Extended Data Fig. 4a-d**). Similar reduction is observed when the length or number of ER sheets is normalized by total cell area (**Extended Data Fig. 4e-i**). Altogether these data demonstrate that the transition in the liver state from physiological to pathological condition in obesity is associated with drastic changes in ER architecture characterized by a disorganization of ER sheets and a transition from ER sheets to tubules.

In liver, ER sheets are often studded with ribosomes forming the rough ER, while the ER tubules are mostly devoid of ribosomes^16^. Given the striking decrease in parallel organized stacks of ER sheets in obese livers observed in our ultrastructure analysis, we evaluated the abundance of the rough and smooth ER. We performed differential centrifugation analysis using discontinuous sucrose gradient and purified rough and smooth ER from liver tissue of lean and obese mice. To validate the efficiency of subcellular fractionation, we also imaged the recovered fractions. As shown in **Extended Data Fig. 5a**, the ER recovered in the denser sucrose fraction is enriched in ribosomes in both lean and obese samples, while the smooth ER fraction was characterized by microsomes vesicles devoid of ribosomes (**Extended Data Fig. 5b**). The amount of rough ER normalized by tissue weight was significantly reduced in livers from obese mice (**Extended Data Fig. 5c**) while the amount of smooth ER tends to be higher (**Extended Data Fig. 5d**). Accordingly, the ratio between rough ER normalized by smooth ER was significantly lower in obese livers (**Extended Data Fig. 5e**).

The different ER subdomains such as rough ER sheets or smooth ER tubules, are shown to be enriched in distinct proteins^17^. ER sheets is the preferential home for the translocon complex of proteins involved in the translocation and modification of newly synthesized polypeptides along with membrane-bound polysomes. Given that the presence of the translocon is dependent on the ribosomes and ER sheets, we evaluated the levels of the translocon channel and its associated proteins. We isolated primary hepatocytes from lean (Wt) and obese (*ob/ob*) mice and stained for the endogenous SEC61α protein. These experiments showed clear SEC61 segregation to the ER sheets in hepatocytes, visualized as patches in the perinuclear ER and cell periphery in hepatocyte of lean mouse (**Extended Data Fig. 5f**). In contrast, the expression of SEC61 is largely decreased and diffused through the whole ER in hepatocytes from obese animals. A similar decrease was also detected liver sections from obese mice compared with the lean controls (**Extended Data Fig. 5g**). Next, we evaluated the total SEC61 expression levels in liver lysates from lean and obese animals. SEC61α, SEC61β and the translocon associated protein TRAPα expression were significantly decreased in lysates from obese mice (**Extended Data Fig. 5h**). Notably, the expression levels of Sec61α, Sec61β and TRAPα were not different in the rough ER fraction isolated from livers between experimental groups, indicating that the decreased total levels of these proteins may reflect primarily the loss of ER sheets rather than a decrease in the translocon complex per rough ER unit (**Extended Data Fig. 5i**). Interestingly, SEC61 and TRAPα levels were actually elevated in the smooth ER, indicating that these proteins translocated to the tubular ER from the sheets. Similar profile was found for GRP78, a protein preferentially found in the rough ER in lean mice. Thus, one clear consequence of the loss of ER sheets in hepatocytes from obese mice is the decrease and mislocalization of the SEC61 and its associated proteins.

The shape of the ER sheets and tubules is determined by several factors including the relative abundance of ER membrane shaping proteins. In mammalian cells, the structure of the tubular ER is determined by the Reticulons and REEPs^14,15,18^. The proteins involved in the formation of ER sheets are less understood. It’s known that the intraluminal space formed between the two ER membranes is established by the presence of a protein called <u>C</u>ytoskeleton-<u>L</u>inking <u>M</u>embrane <u>P</u>rotein 63, Climp-63 (or CKAP4)^13,19,20^. This protein contains a single transmembrane segment and a luminal coiled-coil domain that homo-oligomerize in an anti-parallel manner forming intraluminal bridges ~50nm apart in mammalian cells. Other proteins, such as RRBP1 (p180), are also shown to influence or stabilize ER sheets, yet the impact of p180 on ER sheet formation in mammalian cells is unclear^13,21^. In order to determine whether the loss of ER sheets observed in obese livers is related to alterations in the expression of the ER shaping proteins, we determined their expression profile in total liver lysates and in isolated ER microsomes from lean and obese mice. As shown in **Fig. 3a, b,** the expression of p180 was significantly decreased in total liver lysates of obese animals when compared with their lean counterparts. Although no significant differences were observed in Climp-63 expression levels in total lysates (**Fig. 3a**), this protein was significantly decreased in the total ER fraction (**Fig. 3b**). We found that the expression of Climp-63 and p180 are enriched in the rough ER in liver tissue (**Extended Data Fig. 6a**). In contrast, the expression levels of proteins involved in tubular ER formation such as Reticulon4b and Reep5 were increased in obesity both in the total lysates and total ER fraction (**Fig. 3a, b**). These proteins were preferentially found in our smooth ER fractions (**Extended Data Fig. 6a**). All the antibodies used for these experiments were validated using either positive (overexpression) or negative (deletion) controls (**Extended Data Fig. 6b-f).** We also profiled the mRNA expression levels of multiple genes that encode ER membrane shaping proteins. The increased levels of REEPs and Reticulon 4 were paralleled by increased mRNA expression levels in liver tissues of obese mice (**Extended Data Fig. 6g**). Interestingly, and unlike its protein, Climp-63 mRNA level was higher in the obese samples (**Extended Data Fig. 6g**), possibly indicating a compensatory regulation. Thus, the altered regulation of the abundance of ER shaping proteins promoted by obesity is likely a mechanism through which obesity shift the balance of ER sheet/tubule ratio.

**Fig. 3.**
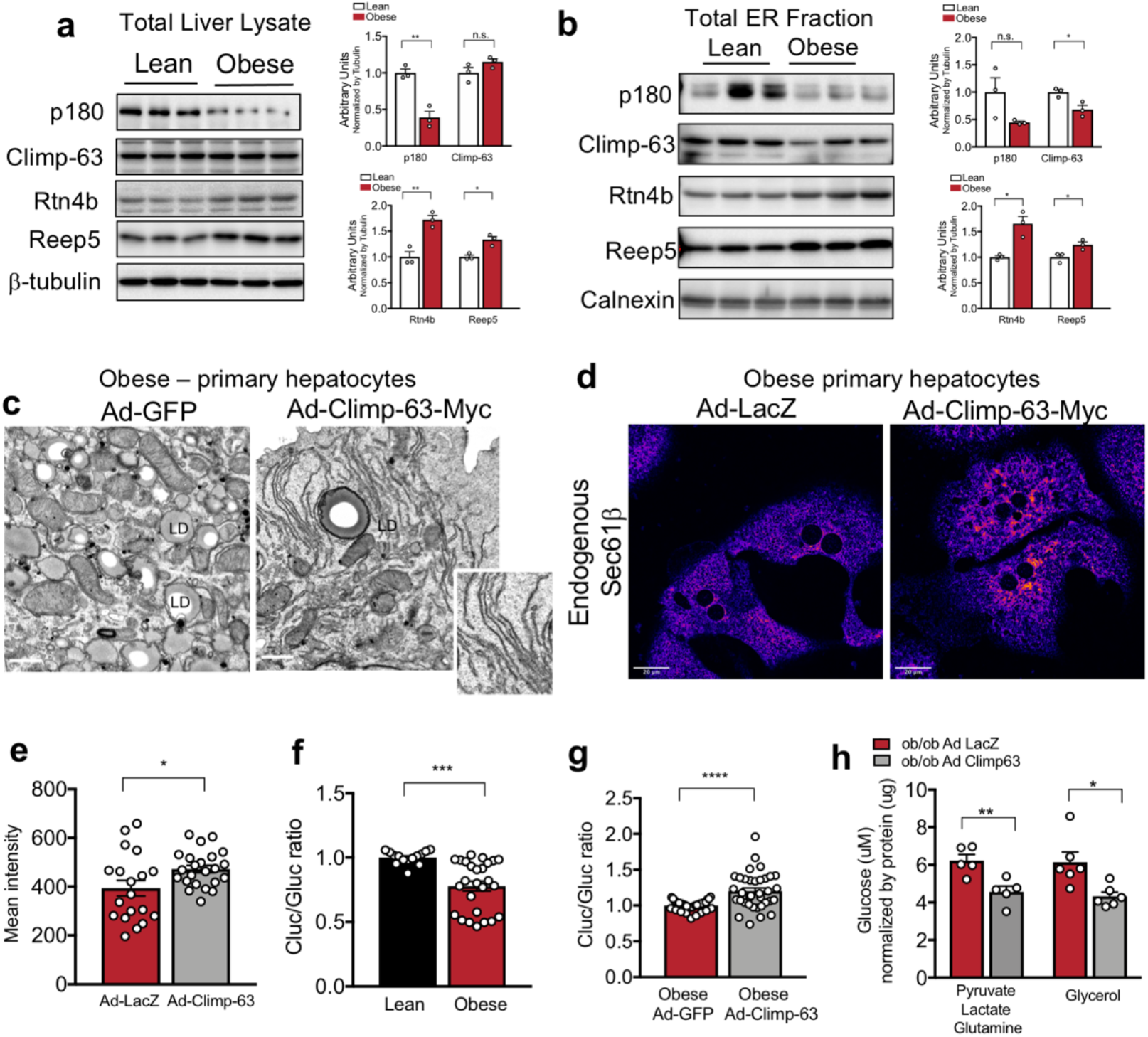
ER shaping proteins are regulated with obesity and exogenous expression of Climp-63 improves function of hepatocytes from obese mice. **a,** Left panel: Immunoblot analysis of indicated proteins in total liver lysates. Right panel: Quantification of the immunoblots. n=3 for lean and obese (*p<0.03, **p<0.005). **b,** Left panel: Immunoblot analysis of indicated proteins from ER fraction isolated from livers from lean and obese. Right panel: Quantification of the immunoblots. n=3 for lean and obese (*p<0.025, **p=0.001). **c,** Representative TEM from primary hepatocytes derived from obese mice exogenously expressing GFP (left) and Climp-63-Myc (right). **d,** Confocal images of immunofluorescence staining for endogenous Sec61β in primary hepatocytes from obese mice exogenously expressing LacZ (left) and Climp-63-Myc (right). **e,** Quantification of fluorescence signal. n=19 fields for LacZ and n=23 fields for Climp-63 (*p=0.0241), representative of 3 experiments. **f,** Quantification of the ratio between luminescence signal from Cluc normalized by luminescence signal derived from Gluc in lean and obese primary hepatocytes. n=14 for lean and n=26 for obese (***p=0.0002). Pooled data from 3 experiments. **g,** Quantification of the ratio between luminescence signal from Cluc normalized by luminescence signal derived from Gluc in obese primary hepatocytes exogenously expressing GFP or Climp-63. n=36 for GFP and n=35 for Climp-63 (****p<0.0001). Pooled from 6 experiments. **h,** Gluconeogenesis assay in primary hepatocytes isolated from obese mice, expressing LacZ (control) or Climp-63, cells are treated with the indicated gluconeogenic substrates in the presence of glucagon for 3 hours. Representative data of 3 experiments.

The relative ratio of ER sheets and tubules is a consequence of a tug of war between ER shaping proteins. In cultured cells, overexpression of Reticulon-homology-domain containing proteins drive ER tubule formation and decrease the abundance of ER sheets, whereas overexpression of Climp-63 leads to enrichment in ER sheets^13,18^. Given the findings described above, we asked whether manipulating ER shape in order to rescue the ER sheets would impact ER function in hepatocytes from obese mice in vitro and in vivo. For that, we used Climp-63 overexpression as a tool to re-shape the ER towards ER sheet proliferation. To confirm the induction of ER sheet proliferation upon exogenous delivery of Climp-63, we first overexpressed Climp-63 in Cos-7 cells (a monkey kidney derived cell line). Fluorescence images presented in **Extended Data Fig. 7a,** showed that exogenous expression of Climp-63 results in a massive increase in ER sheets (as patches around the nucleus) in Cos-7 cells as shown in previous reports^13^. We also examined the proliferation of ER sheets driven by Climp-63 overexpression by TEM, where we observed a striking proliferation of membrane sheets with a constant luminal space, stacked very closely together **(Extended Data Fig. 7b)**. We then cloned Climp-63 in an adenovirus backbone in order to exogenously express this protein in primary hepatocytes isolated from lean and obese mice. As controls, we used adenoviruses expressing either LacZ or GFP. Exogenous expression of Climp-63 in primary hepatocytes lead to a wide proliferation of ER sheets in primary hepatocytes both in lean (**Extended Data Fig. 7c**) and obese samples as examined by confocal microscopy and TEM (**Fig. 3c**). We then asked whether the increase in ER sheets affect SEC61 expression and localization. We stained endogenous SEC61 in the ER from the hepatocytes. As shown in **Fig. 3d, e**, we detected that overexpression of Climp-63 in primary hepatocytes obtained from obese mice led to a recovery of Sec61β localization to the ER sheets, as shown by endogenous staining for Sec61β in these cells.

We then tested if increasing ER sheets and recovery of translocon localization impacts ER’s protein folding and secretory capacity. For that, we used a luciferase based genetic reporter^22^, based on the folding and secretion of the membrane protein asialoglycoprotein receptor 1 (ASGR1), a slow-folding protein whose secretion is decreased under ER stress conditions^22^ (construct described in **Extended Data Fig. 7e**). To evaluate the impact of structural changes on ER folding and secretory capacity in hepatocytes, we infected primary hepatocytes isolated from lean and obese animals with an adenovirus expressing the ASGR1 reporter. As shown in **Fig. 3f,** the secretion of ASGR-Cluc is decreased in primary hepatocytes derived from obese animals compared with control primary hepatocytes. Exogenous expression of Climp-63 in hepatocytes from obese animals lead to increased folding capacity compared with obese cells expressing adenovirus LacZ control (**Fig. 3g**).

We then evaluated the impact of ER re-shaping towards increased sheet formation in obese hepatocytes on another metabolic function, glucose production. As shown in **Fig. 3h,** exogenous expression of Climp-63 leads to decreased glucose production in primary hepatocytes from obese mice, driven by different gluconeogenic substrates. Thus, correcting the architectural organization of the ER in livers from obese mice is sufficient to improve hepatic ER proteostasis and metabolic function and demonstrates the ER shape directedly affect hepatocyte metabolic program.

Given the promising results in vitro, we tested whether structural manipulation through Climp-63 overexpression could impact liver and systemic metabolism in vivo. For that, we delivered LacZ (control) or Climp-63 to livers of 10 weeks old lean and obese mice with adenoviral vectors. As shown in **Fig. 4a, b, and Extended Data Fig. 7f,** overexpression of Climp-63 in vivo led to a significant increase in parallel ER sheet formation both in lean and obese mice livers. Noticeably, the sheets promoted by Climp-63 overexpression in liver tissue were decorated with ribosomes as endogenous ER sheets (**Fig. 4a *inset***). It’s interesting to note that the presence of ribosomes in exogenously induced ER sheet formation was not observed in Cos7 cells (**Extended Data Fig. 7b**). This illustrates the critical importance of native context in order to establish structure/function relationship.

**Fig. 4.**
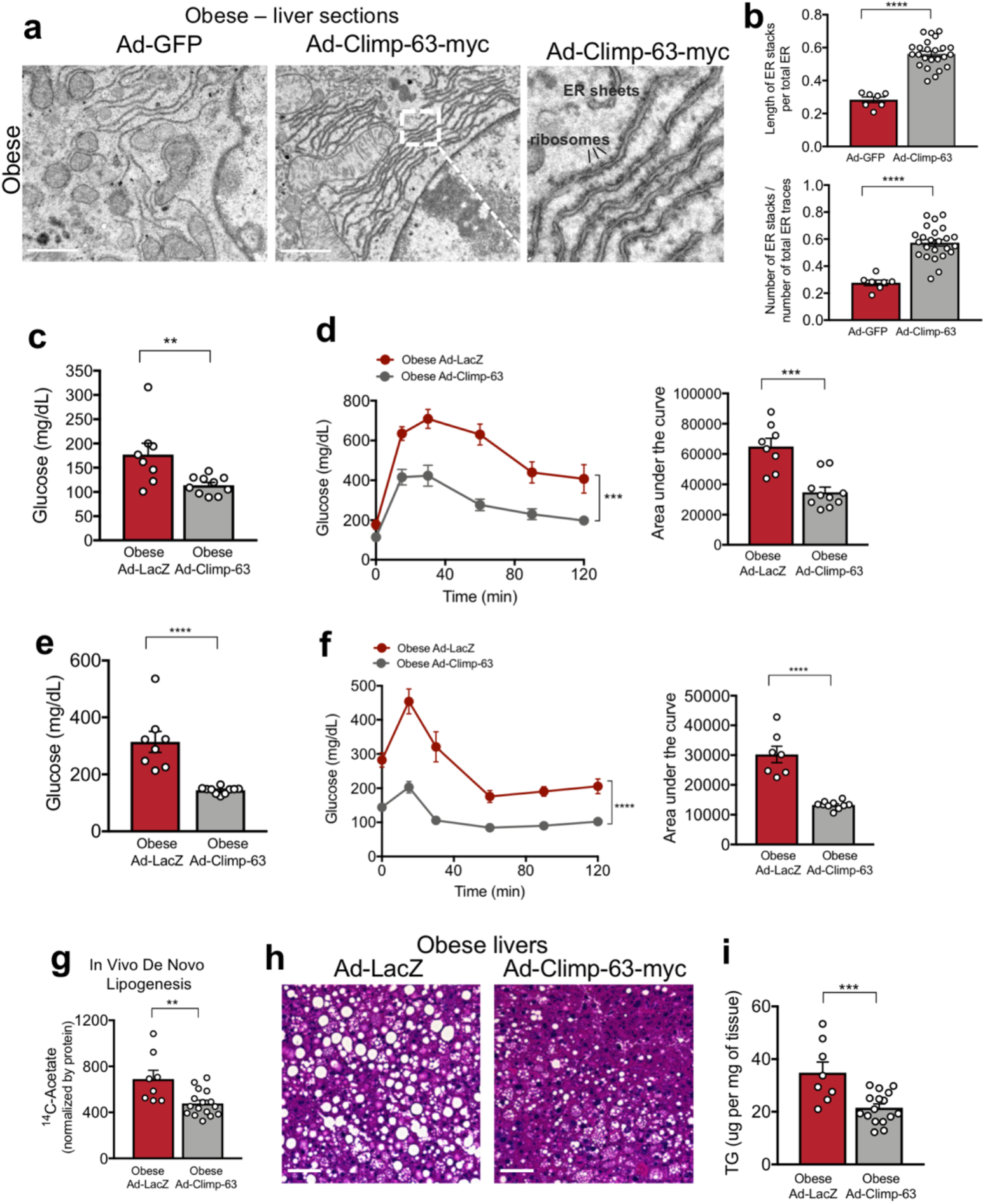
Exogenous expression of Climp-63 in livers of obese mice improves systemic glucose and lipid metabolism. **a,** Representative TEM from livers derived from obese mice exogenously expressing GFP (left) and Climp-63-Myc (right) Ad: Adenovirus. Scale bar: 1.118um. Inset: High magnification image of ribosomes associated with ER sheets promoted by Climp-63 overexpression. **b,** Quantification of parallel organized stacks of ER in liver sections presented in figure 5A. n=7 for GFP and n=24 Climp-63-myc EM sections (****p<0.0001). **c,** Blood glucose levels from 11 weeks old obese mice exogenously expressing LacZ or Climp-63 after overnight fasting. n=8 for LacZ and n=10 for Climp-63-myc. Representative of 3 independent cohorts (**p=0.0098). **d,** Left: Glucose tolerance test in obese mice exogenously expressing LacZ (control) and Climp-63-myc. Right panel: area under the curve of the experiment in left panel. n=8 ad-LacZ and n=10 ad-Climp-63-myc animals, representative of 3 independent experiments (***p=0.0002). **e,** Blood glucose levels from 10 weeks old obese mice exogenously expressing LacZ and Climp-63-myc after 6 hours of food withdrawn. n=8 for LacZ and n=10 for Climp-63-myc. Representative of 3 independent experiments (****p<0.0001). **f,** Insulin tolerance test in obese mice exogenously expressing LacZ (control) and Climp-63-myc. n=8 for LacZ and n=10 for Climp-63-myc. Representative of 3 independent experiments (****p<0.0001). **g,** ^14^C-Acetate driven de novo lipogenesis assay; scintillation counts per minute normalized by mg of liver tissue. n=8 liver lobes from 2 obese mice expressing LacZ and n=16 liver lobes from 4 obese mice expressing Climp-63 exogenously (***p=0.0010). **h,** Histological analysis of liver sections derived from obese mice exogenously expressing GFP (left) and Climp-63 (right) stained with hematoxylin and Eosin (H&E). Scale bar: 65.4 um. **i,** Triglyceride content of the same livers in Fig. 4c (***p=0.0037).

We next assessed the effect of exogenous expression of Climp-63 in liver tissues of obese mice on glucose metabolism. As shown in **Fig. 4c**, expression of Climp-63 in obese mice resulted in a significant decrease in blood glucose levels after 16h fasting and a striking improvement in glucose clearance evaluated by a glucose tolerance test (**Fig. 4d**). Blood glucose levels were also markedly lower in obese mice overexpressing Climp-63 compared to the controls expressing LacZ in the fed state (**Fig. 4e**). Insulin sensitivity was also significantly improved by Climp-63 overexpression (**Fig. 4f**). We next examined the impact of recovering ER sheets in obesity on the rate of de novo lipid synthesis in the liver. As shown in **Fig. 4g** exogenous expression of Climp-63 led to decreased hepatocyte lipogenic capacity measured through the incorporation of ^14^C-labeled-acetate in de novo lipogenesis assay in vivo. We then asked whether this reduction in lipogenic activity is reflected in liver lipid accumulation. In agreement with the reduced *de novo lipogenesis*, we observed that Climp-63 overexpression led to a significant decrease in lipid accumulation in the liver by both histological examination (**Fig. 4h**) and total triglyceride content in the liver (**Fig. 4i**). Thus, rescuing ER sheet/tubule ratio had a direct impact in liver lipid synthesis and steatosis in the liver. Altogether these data show that correcting the architectural organization of the ER in livers from obese mice exerts significant effects on hepatic and systemic metabolism. Hence, by simply changing ER structure, it is possible to alter hepatocyte function and metabolic homeostasis revealing the essential relationship between organelle structure and function.

In fact, in all systems and at all levels, structure is a critical determinant of function. In static systems, only a limited number of tasks can be achieved within the constraints of a rigid and inflexible construction. In living cells, the vast diversity and highly dynamic nature of tasks demand sub-cellular structural complexity as well as flexibility to support functional integrity and survival and to maximize the repertoire of proper and compartmentalized responses generated from biological infrastructure. Metabolic processes are also exceedingly complex and compartmentalized and demand high level of adaptive flexibility. Here, we used the ER as one of the central architectural assembly of the cells and provide what we believe to be, the most detailed characterization of the subcellular architecture in liver tissue in both health and disease context. This analysis included the precise visualization and quantification of hepatic tubular ER ultrastructure, a complex structure which has been very challenging to capture in detail in tissue setting with other imaging approaches. Additionally, we examined structure-function relationship in the context of metabolic homeostasis in health and disease, using obesity as a model. The impact of the increased ER sheets on metabolism may occur as a consequence of the activity and abundance of proteins/enzymes that are housed in this ER subdomain or maybe downstream of hormonal or metabolic signals that govern metabolic output. For example, we showed that rescuing ER sheets reflected in the recovery of the Sec61 translocon complex and polysome association with ER membrane which likely impact ER folding capacity as shown in **Fig. 3g**. It is also possible that Climp-63 overexpression alters ER interactions with microtubules^23–26^ and restricts the lateral mobility of translocon complex^16,24,27–29^. Regardless, an exciting prospect of our observations is that structure is a prerequisite for metabolic programming, and such a regulatory circuit could open up new avenues in understanding endocrine and metabolic homeostasis including response to hormonal or nutritional cues to determine the metabolic outcomes.

The data generated by this work provide the field a significant amount of information that can be further explored with different questions related to liver subcellular architecture. It’s important to highlight that this analysis involved extensive sample preparation, high resolution imaging, automated image segmentation and reconstruction. While all of which were extremely demanding processes in terms of human labor and computing power, in our view, this workflow allows precise understanding of the structure-function relationship in the native tissue context and how this could be possibly linked to metabolic function in health and disease and exploited for diverse therapeutic opportunities.

---

**Extended Data Fig. 1.**
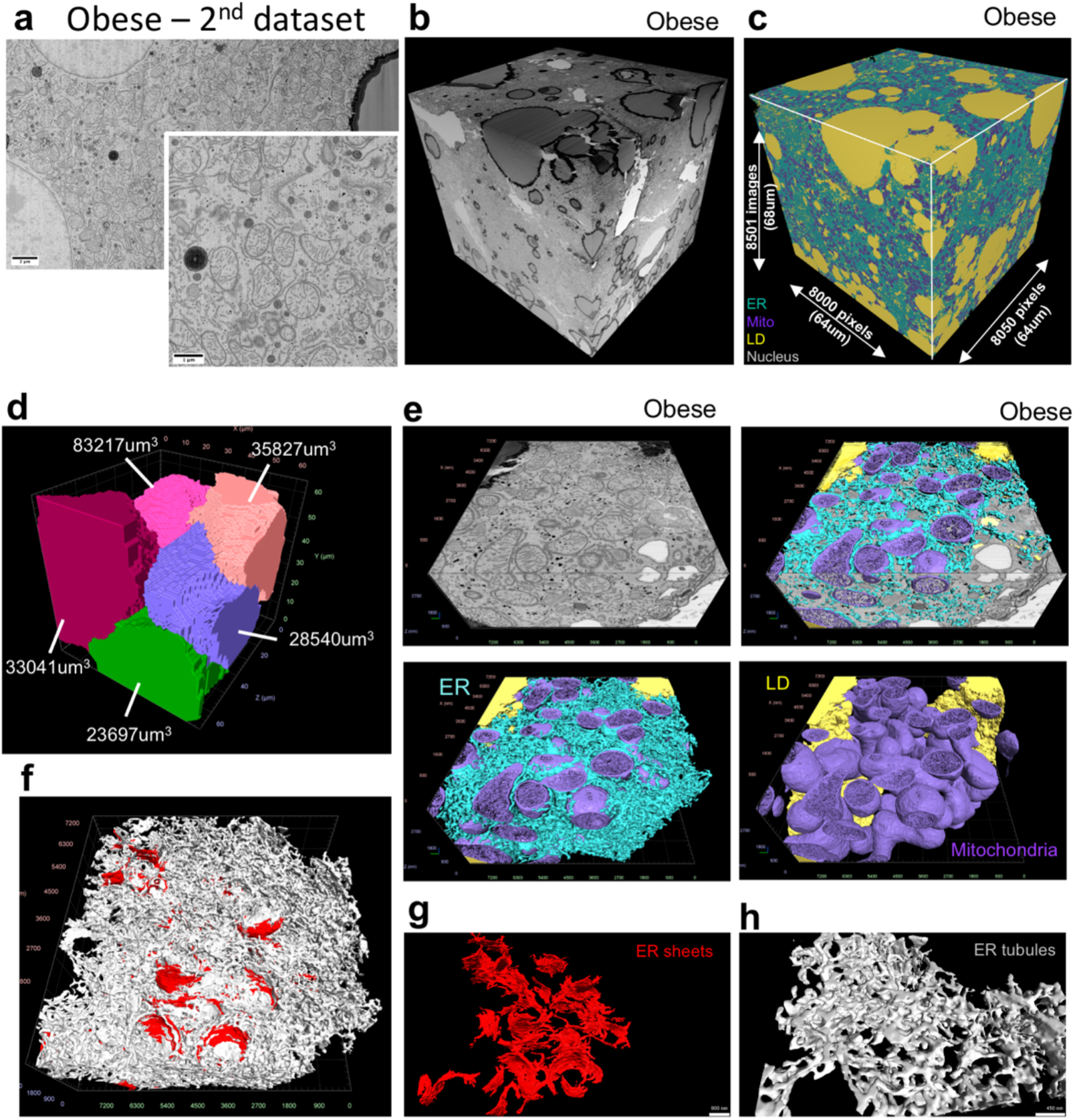
FIB-SEM imaging and automated deep-learning-based segmentation of the 2^nd^ dataset from obese liver. **a,** Single section SEM of liver from obese (*2*^*nd*^ *dataset*) mice liver in fed state at 8 nm pixel size. **b,** 3D reconstruction of FIB-SEM images derived from obese liver volume. **c,** Convolutional neural network based automated segmentation of liver volumes derived from obese mice. The dimensions of the volume are depicted in the figure. ER (endoplasmic reticulum, blue), Mito (mitochondria, purple), LD (lipid droplet, yellow), Nucleus (gray). **d,** Reconstruction of 5 full or partial hepatocyte volumes present in the liver volume imaged by FIB-SEM. The volumes of the cells are depicted in the figure. All the reconstructions were performed in Arivis Vision 4D software. **e,** Section of the liver from obese mice segmented with ER (blue), mitochondria (purple), cristae (pink) and lipid droplet (yellow) annotation and reconstruction. **f,** Sub-segmentation and 3D reconstruction of ER sheets (red) and tubules (gray) from obese mice. **g,** Example volumes of ER sheets and **h,** ER tubules from this dataset.

**Extended Data Fig. 2.**
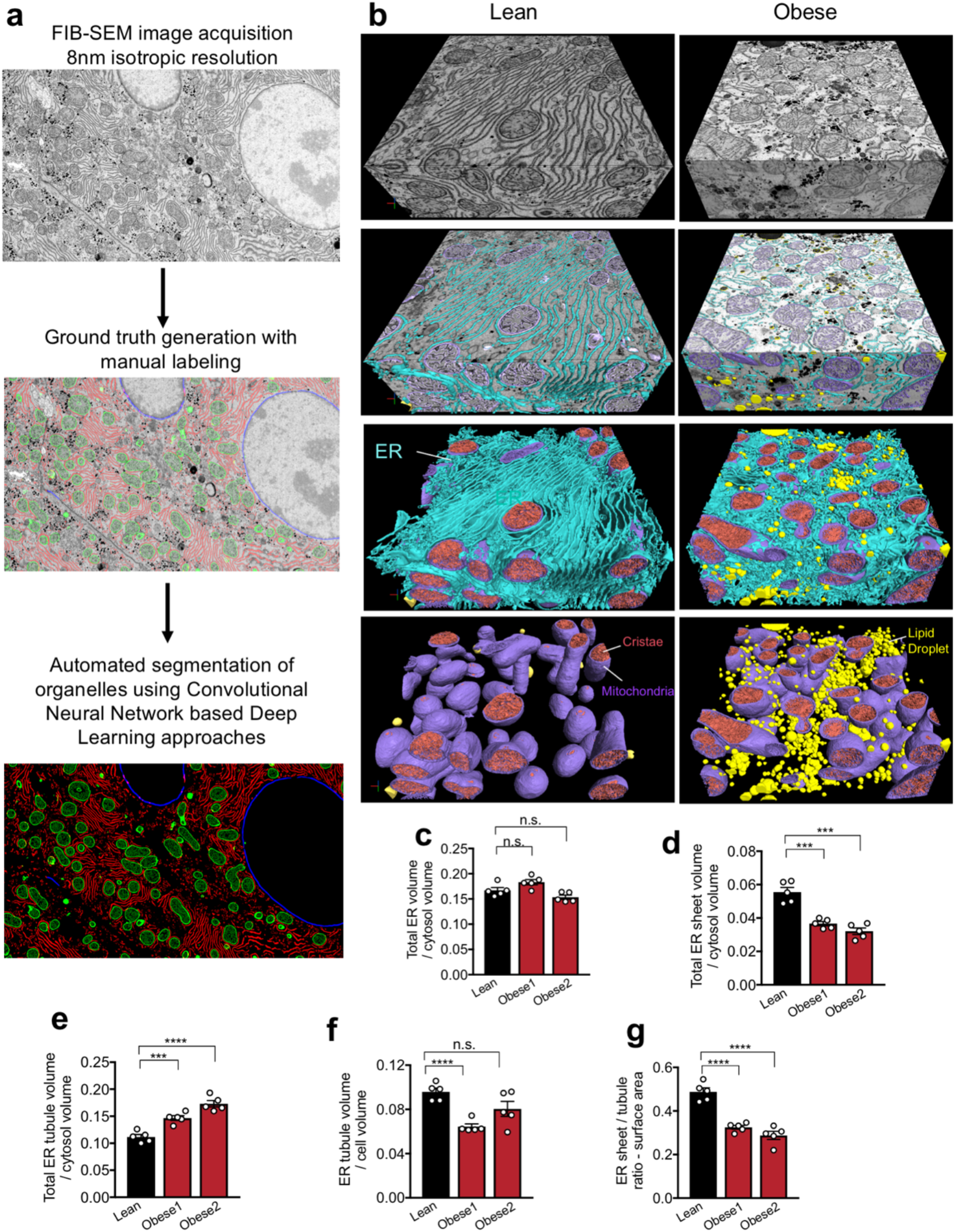
Workflow of automated deep-learning-based segmentation. **a,** Workflow for automated segmentation of organelles using convolution neuronal network-based machine learning. **b,** Section of the liver volume from lean (right) and obese (left) with ER (blue), mitochondria (purple), cristae (pink) and lipid droplet (yellow) annotation and reconstruction. **c,** Quantification of total ER volume area normalized by cytosol volume (here cytosol was considered the cell volume minus the volume occupied by lipid droplets, mitochondria and ER). n=5 for each group. **d,** Quantification of ER sheet volume normalized by cytosol volume. n=5 per group (****p<0.0001). **e,** Quantification of ER tubule volume normalized by cytosol volume. n=5 per group (***p=0.0006, ****p<0.0001). **f,** Quantification of ER tubule volume area normalized by cell volume. n=5 for each group (****p<0.0001). **g,** Quantification of total ER surface area normalized by cell surface area in 5 different cells derived from lean and obese liver (****p<0.0001).

**Extended Data Fig. 3.**
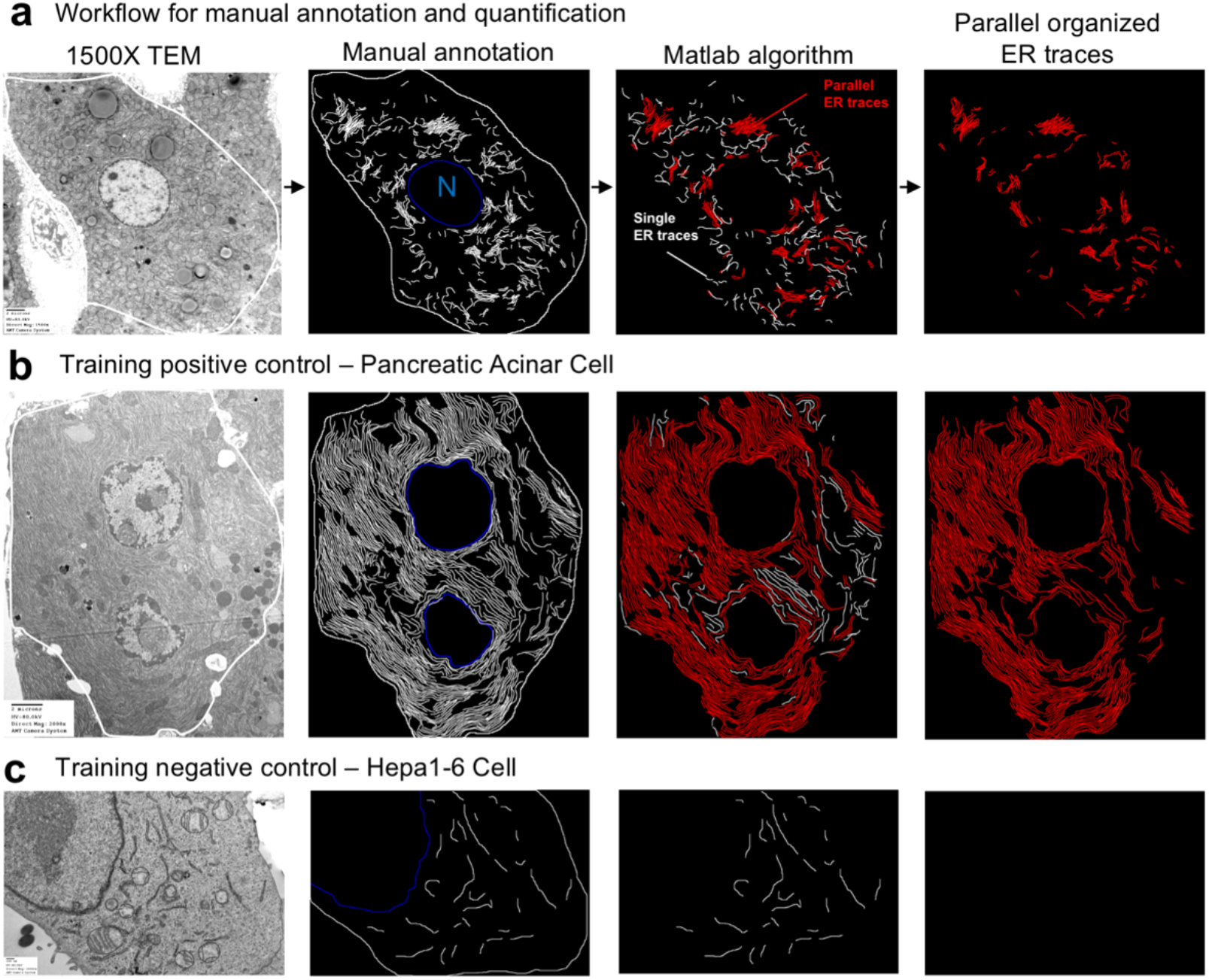
MATLAB algorithm to quantify parallel organization of ER membrane. **a,** Workflow for manual annotation and quantification of parallel organized ER sheets. We considered two neighboring ER sheets as “parallel” if more than 50% of the two neighboring ER traces are within in 55-225 nm distance range (5-20 pixel range) from each other. **b,** TEM of acinar cell section used as a positive control for training the algorithm. **c,** TEM of Hepa 1-6 cell used as a negative control for training the algorithm.

**Extended Data Fig. 4.**
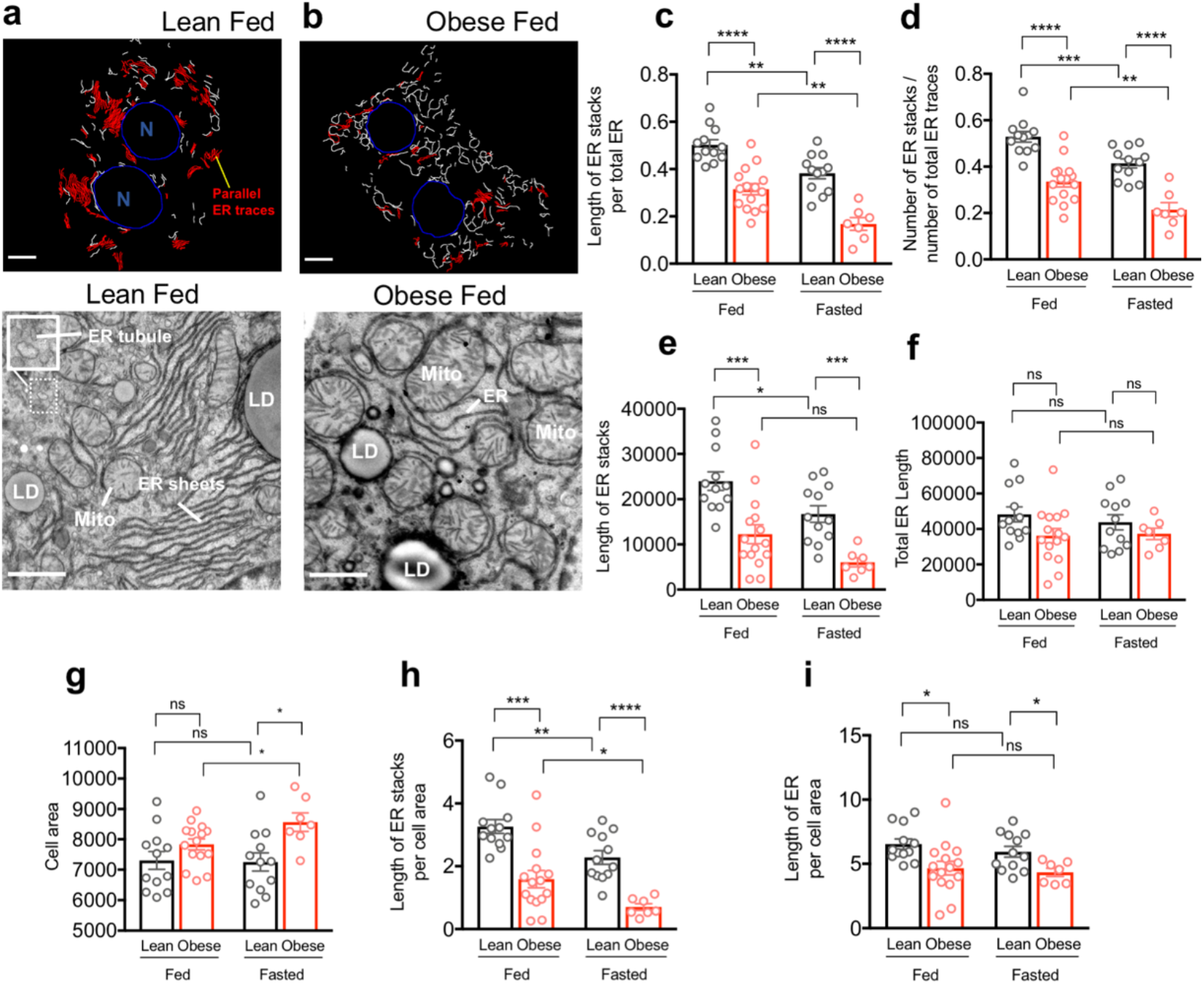
Parallel organized stacks of ER sheets are decreased in obesity. **a, b,** Binary masks of manually annotated ER from the TEM images acquired in 1500x mag. White and red represent ER traces, where parallel organized ER is segmented in red, blue: nucleus (N). Bottom images are representative transmission electron microscopy (TEM) images of liver sections derived from lean and obese mice in fed state. ER (endoplasmic reticulum), Mito (mitochondria), LD (lipid droplet). Scale bar: 2.236um. **c,** Quantification of the length of parallel ER stacks normalized by total ER length (****p<0.0001, **p<0.0011). **d,** number of parallel ER stacks normalized by total number of ER traces (****p<0.0001, ***p=0.0010, **p=0.0067). **e,** Quantification of the length of total parallel ER sheets traces (***p=0.0006, *p=0.015). **f,** Quantification of the total ER length. **g,** Quantification of total cell area (*p=<0.04). **h,** Quantification of the length of parallel ER stacks normalized by cell area (***p=<0.0001, **p=0.0043, *p=0.0457). **i,** Quantification of length of total ER traces normalized by cell area.

**Extended Data Fig. 5.**
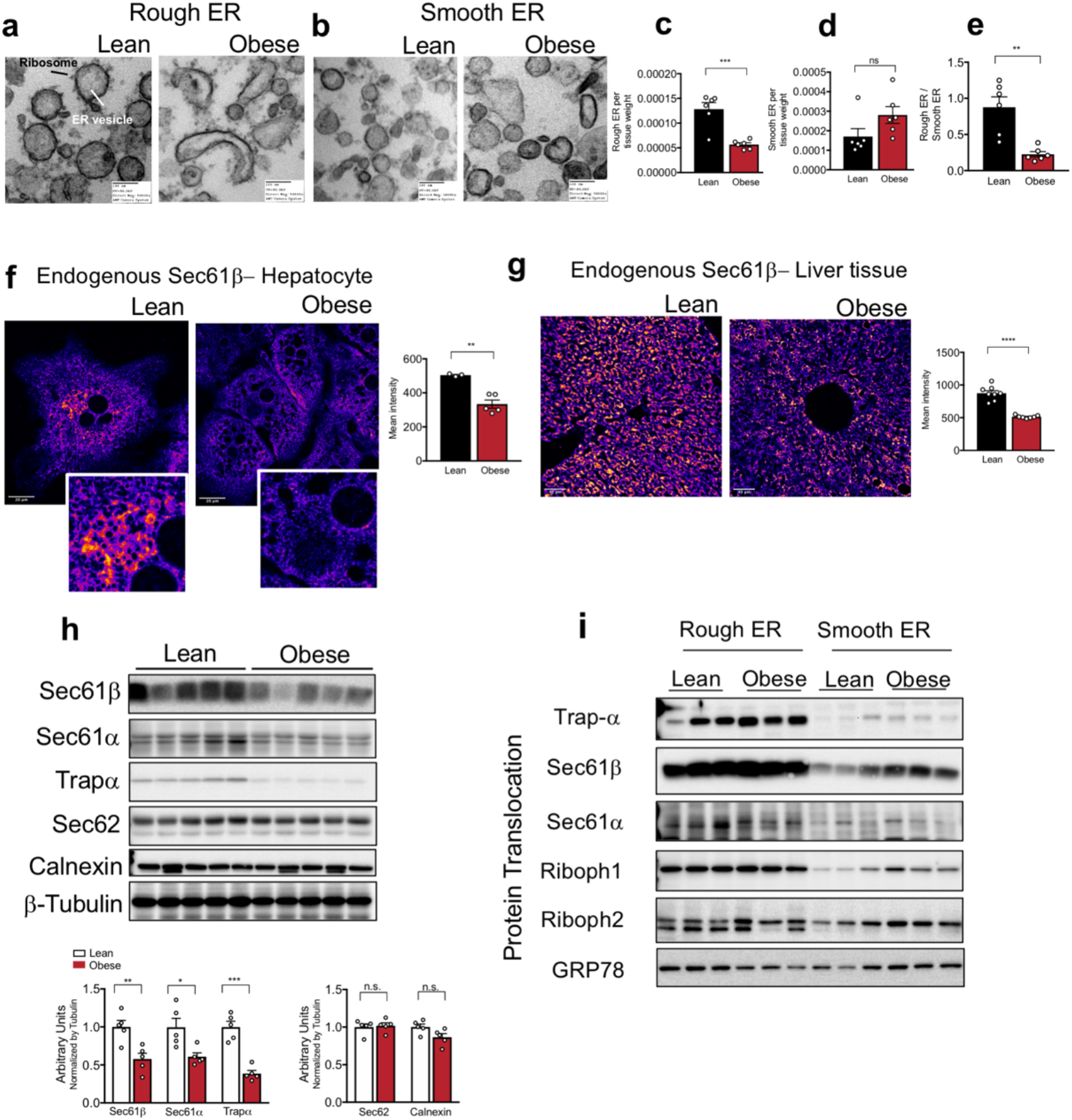
Rough ER and translocon complex are downregulated in obesity. **a,** TEM of rough and **b,** smooth ER vesicles isolated from livers derived from lean and obese mice. **c,** milligrams of rough (***p=0.0004) and **d,** smooth (p=0.0917) ER vesicles recovered by subcellular fractionation normalized by milligram of liver. **e,** Ratio between abundance of rough divided by smooth ER vesicles (**p=0.0014). The graphs represent average of 6 mice for lean and 6 mice for obese. **f,** Confocal images and quantification of immunofluorescence staining for endogenous Sec61β in primary hepatocytes from lean and obese mice. n=3 fields lean and n=5 fields obese, representative of 3 independent hepatocytes isolations (**p=0.0018). **g,** Confocal images of immunofluorescence staining for endogenous Sec61β in liver sections from lean and obese mice. Right panel: Quantification of fluorescence intensity of immunofluorescence staining for endogenous Sec61β in liver sections from lean and obese mice. n=8 fields for lean and 7 fields for obese mice, representative of 2 mice per group (****p=0.0001). **h,** Upper panel: Immunoblot analysis of the indicated proteins in total liver lysates from lean and obese mice. Bottom panel: Quantification of the immunoblots. n=5 mice per group, representative of 3 independent cohorts. In all panels error bars denote s.e.m. **i,** Immunoblot analysis of indicated proteins in rough and smooth ER fractions from liver tissue derived from lean and obese mice.

**Extended Data Fig. 6.**
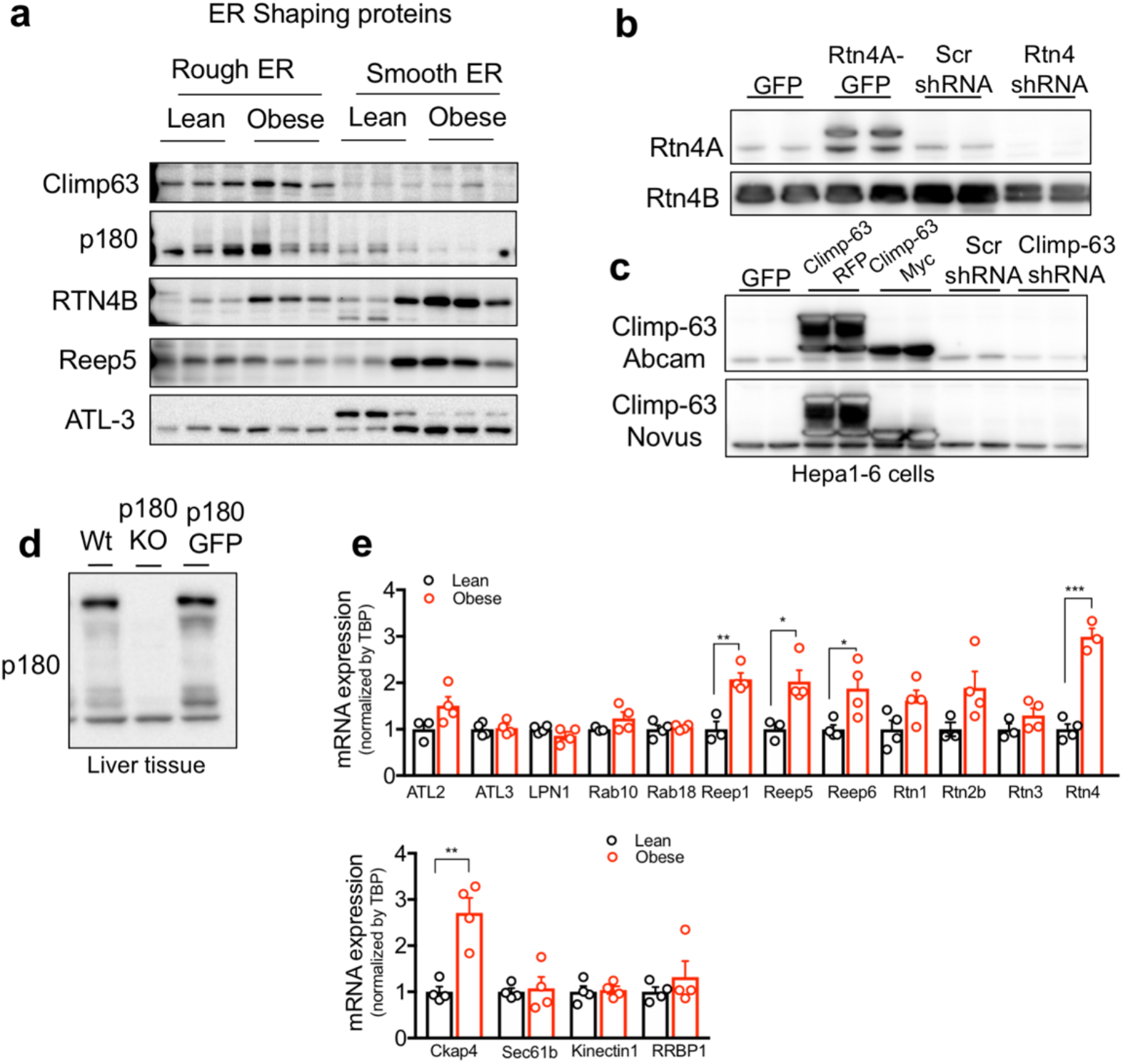
Analysis of shaping proteins. **a,** Immunoblot analysis of indicated proteins in rough and smooth ER fractions from livers derived from lean and obese mice. **b,** Immunoblot analysis for Reticulon 4A and 4B in total lysates from Hepa1-6 cells overexpressing Rtn4A tagged with GFP and Hepa1-6 cells transfected with shRNA control (scrambled, Scr) and shRNA against Reticulon4. **c,** Immunoblot analysis for Climp-63 in total lysates from Hepa1-6 cells expressing GFP control and Climp-63 tagged with RFP or Myc and Hepa 1-6 cells expressing shRNA control (scrambled, Scr) and shRNA against Climp-63. **d,** Immunoblot analysis for RRBP1 (p180) in total liver lysates derived from wild type (Wt) and RRBP1 deficient mice. **e,** mRNA expression levels for indicated genes measured by qPCR. cDNA samples derived from lean and obese mice livers. n=4 for lean and 4 for obese (*p<0.027, **p<0.004, ***p=0.0002).

**Extended Data Fig. 7.**
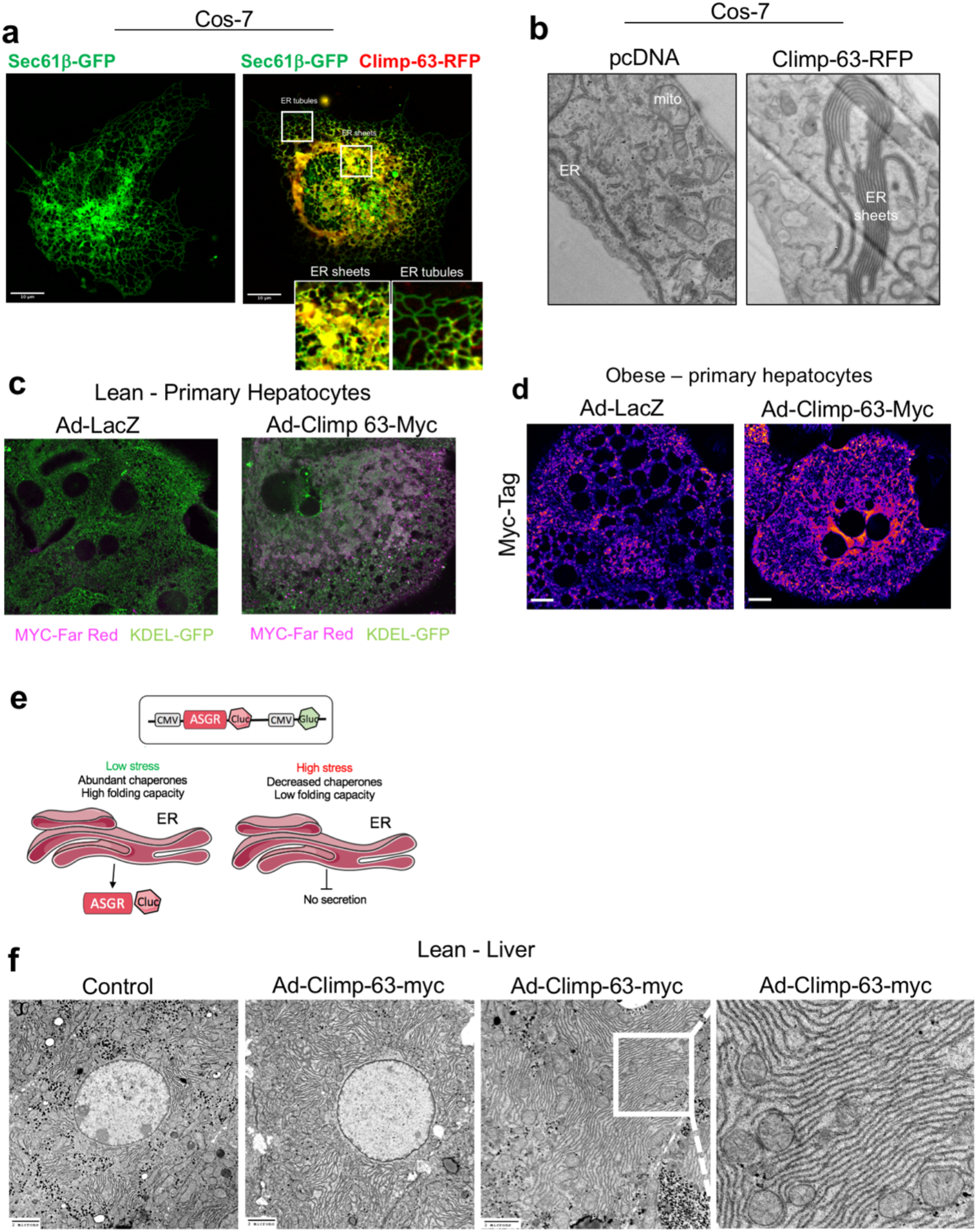
Exogenous expression of Climp-63 in primary hepatocytes and in liver in vivo promotes ER sheet formation. **a,** Left panel: Confocal images from Cos-7 cells exogenously expressing Sec61β fused with GFP as a fluorescent general ER marker. Right panel: Overlay of images from Cos-7 cells exogenously expressing Sec61β fused with GFP (green) and Climp-63 fused with RFP (red). Overlay is shown in yellow. **b,** Representative TEM from Cos-7 cell sections expressing control (pcDNA) or Climp-63-RFP constructs. **c,** Endogenous staining of KDEL as an ER marker (in green) and Myc-tag (in far red) in lean primary hepatocytes expressing either Ad-LacZ or Ad-Climp-63-Myc. **d,** Confocal images of immunofluorescence staining for Myc reflecting Climp-63-Myc expression in primary hepatocytes from obese mice, expressing LacZ or Climp-63-Myc Ad (adenovirus). Scale bar: 10.96 um. Signal on the left image is un-specific staining from the anti-Myc antibody. **e,** Scheme describing the ASGR reporter. **f,** Representative TEM from liver sections derived from lean control mice or lean mice overexpressing Climp-63-Myc in vivo.

## Supplementary videos can be reached at

https://www.youtube.com/playlist?list=PLpzquMkvsJ9XZvqAKyHcz62iDQkoKxmpx

## Supplementary video Legends

**Supplementary video 1. 3D reconstruction and segmentation of FIB-SEM images derived from liver volume from lean mice.** Reconstruction of 5638 consecutive images of a liver tissue volume corresponding to 96μm(x), 64μm(y), 45μm(z) at voxel size of 8 nm in x, y, and z dimensions. ER is segmented in green, mitochondria in purple and lipid droplets in yellow. (**Fig. 1c**).

**Supplementary video 2. 3D reconstruction and segmentation of FIB-SEM images derived from liver volume from obese mice.** Reconstruction of 7896 consecutive images of a liver tissue volume corresponding to 81μm(x), 73μm(y), 63μm(z) at voxel size of 8 nm in x, y, and z dimensions. ER is segmented in green, mitochondria in purple and lipid droplets in yellow. (**Fig. 1d**).

**Supplementary video 3. Segmentation of organelles in liver volume from lean mice.** ER (green), mitochondria (purple), lipid droplets (yellow) and nucleus (gray), in a volume of 5.4×10^11^ voxels of liver tissue, were segmented using deep learning - convolutional neural networks (CNN). (**Fig. 1e**).

**Supplementary video 4. Segmentation of organelles in liver volume from obese mice.** ER (green), mitochondria (purple), lipid droplets (yellow) and nucleus (gray), in a volume of 7.3×10^11^ voxels of liver tissue, were segmented using deep learning - convolutional neural networks (CNN). (**Fig. 1f**).

**Supplementary video 5. Visualization of ER organization in sub-volume of a hepatocyte from lean mice.** ER (green) and mitochondria (purple) highlighting the predominance of ER sheets in the sample (1000×1000×400 pixels). (**Fig. 2a**).

**Supplementary video 6. Visualization of ER organization in sub-volume of a hepatocyte from obese mice.** ER (green) and mitochondria (purple) highlighting the predominance of ER tubules in the sample (1000×1000×400 pixels) (**Fig. 2b**).

**Supplementary video 7. Fly-through visualization of ER and mitochondria organization in liver volume from lean mice.** A liver volume of 8×8×3.2 um^3^ was rendered using Houdini SideFX. Detailed views of ER sheets and tubules are shown in white and mitochondria and cristae are shown in red. (**Fig. 2c**).

**Supplementary video 8 and 9. Fly-through visualization of ER and mitochondria organization in liver volume from obese mice.** Liver volumes of 8×8×3.2 um^3^ from two separate obese mice were rendered using Houdini SideFX. Detailed views of ER sheets and tubules are shown in white and mitochondria and cristae are shown in red. (**Fig. 2d and Extended Data Fig. 1**).

**Supplementary Table 1:** Dataset volumes.

| Dataset (Liver) | X (pixels) | Y (pixels) | Z (pixels) | Segmented volume size ($\mu\text{m}^3$) | Number of classes segmented (raw included) | Data Size for 1 class (Tiff files) | Total Data Size |
| --- | --- | --- | --- | --- | --- | --- | --- |
| Lean | 12000 | 8000 | 5638 | 277119 | 9 | 541 Gb | 4.87 Tb |
| Obese1 | 9112 | 10200 | 7896 | 375743 | 10 | 734 Gb | 7.34 Tb |
| Obese2 | 8000 | 8050 | 8501 | 282633 | 7 | 548 Gb | 3.84 Tb |

**Classes segmented:** ER, mitochondria, cristae, lipid droplet, Golgi, peroxisome, glycogen, nucleus, plasma membrane

**Supplementary Table 2:**
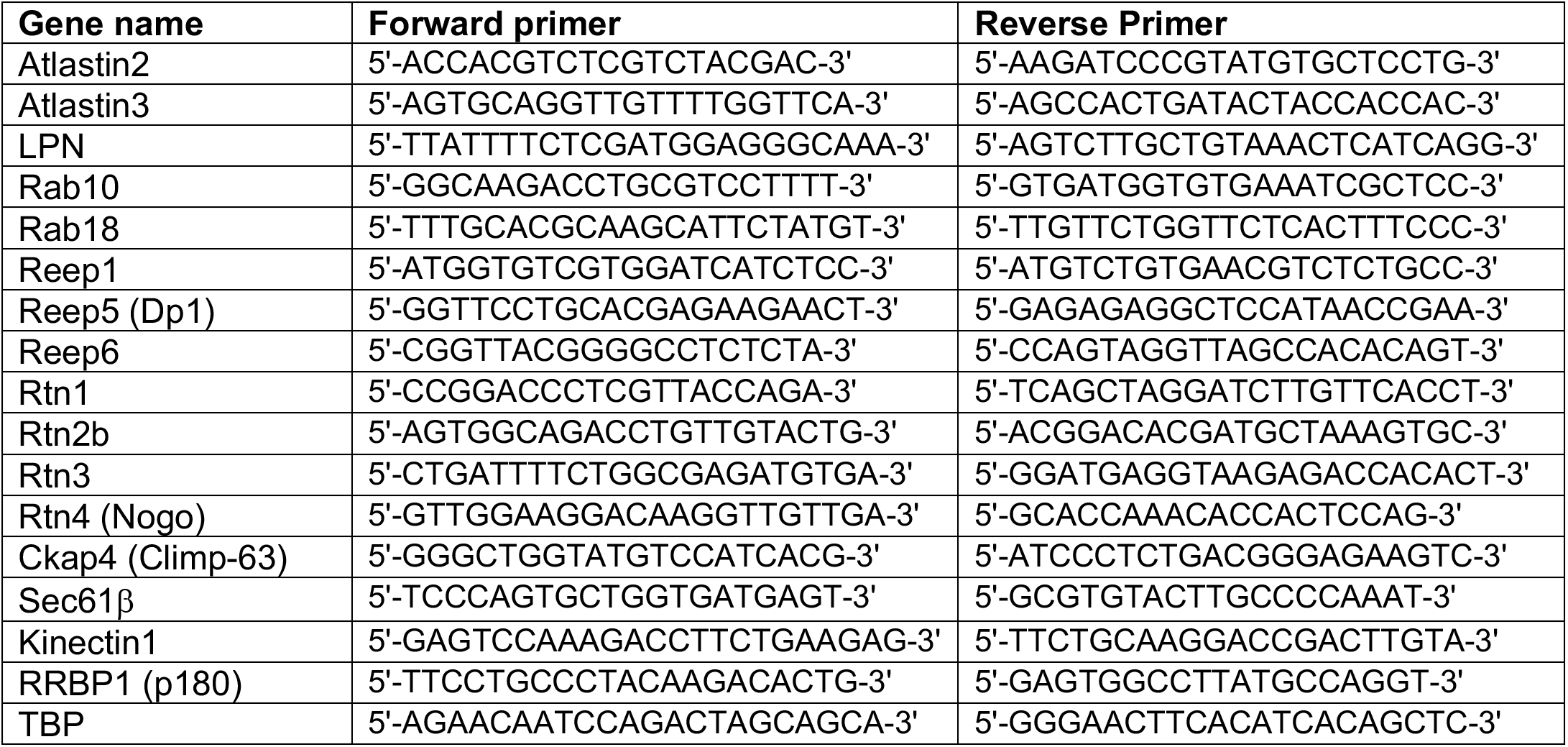
Primer list. (all Mus musculus)

## METHODS

### General animal care, study design and animal models

All experimental procedures involving animals were approved by the Harvard T.H. Chan School of Public Health (HSPH) Institutional Animal Care and Use Committee. The mice were maintained from 6 to 12 weeks old on a 12-hour-light /12-hour-dark cycle in the Harvard T.H. Chan School of Public Health pathogen-free barrier facility with free access to water and to a standard laboratory chow diet (PicoLab Mouse Diet 20 #5058, LabDiet). As an animal model of obesity, we used the leptin deficient *B6.Cg-Lep*^*ob*^/*J* (*ob*/*ob*) mouse in the C57BL/6J genetic background (Stock no. 000632). As controls, we used aged and gender matched ob/+ hets (Stock no. 000632) and homozygous (+/+) (Stock no. 000664). These animals were purchased from Jackson Laboratories at 6-7 weeks of age and used for experimentation between 8-11 weeks of age. In animal experiments, all measurements were included in the analysis. The sample size and number of replicates for this study were chosen based on previous experiments performed in our lab and others^1,2^. No specific power analysis was used to estimate sample size.

### Adenovirus-mediated overexpression of Climp-63

For exogenous Climp-63 (CKAP4) expression, mouse Climp-63 tagged with Myc-DDK in the C terminal (Origene Catalog: MR215622) was cloned in an Ad5 backbone from Vector BioLabs. The adenovirus (serotype 5, Ad5) was generated, amplified and double purified in CsCl by Vector Biolabs. The adenovirus was administered to 8 weeks-old *ob/ob* mice, at a titer of 1.2×10^9^ IFU/mouse. Metabolic studies were performed between day 7-12 after infection. The animals were sacrificed after 14 days of infection for ex vivo experiments.

### Glucose and insulin tolerance tests

For glucose tolerance test, animals were overnight fasted and subjected to an intraperitoneal (i.p.) glucose injection (Wt: 1.5 g kg^−1^, ob/ob: 1.0 g kg^−1^). Blood glucose levels were measured throughout the first 120 minutes of the glucose injection. For the insulin tolerance test, food was removed for 6 hours (from 9 am to 3 pm). Animals were subjected to an i.p. injection of insulin (Wt: 0.8 U/Kg and ob/ob 1.5U/Kg). Insulin was diluted and prepared in 0.1% BSA.

### Liver triglyceride measurements

Liver tissues (~100mg) were homogenized in 1.2mL of water using a TissueLyser (Qiagen) for 5min, at 30f/s, 2 cycles at 4°C. Next, samples were vortexed and 100μL of the homogenate was transferred to a 1.5mL tube and 125μL of chloroform and 250μL of Methanol were added. Samples were vortexed briefly and incubated for 5 min. Additional 125μL of chloroform was added. Next, 125μL of water was added and samples were vortexed and centrifuged at 3000rpm for 20 min at 4°C. Around 150μL of the lower phase was collected in a 1.5mL tube and chloroform was evaporated in a heated vacuum oven. Lipids were then re-suspended in 300uL of ethanol. Triglyceride was measured by a Randox Tg Kit (catalog number TR213) from Randox Laboratories.

### Lipogenesis assay

Obese (ob/ob) mice (~10 weeks old) in the fed state were injected with 1.5U/kg insulin. After 60 minutes, mice were injected with 10μM cold Acetate and 25μCi ^14^C-Acetate diluted in PBS in a total of 250uL. After 60 minutes, animals were sacrificed, total liver was weighted, and 4 pieces of different lobes were collected from each animal and frozen in liquid nitrogen. Lipids were extracted similarly described in Liver Tg Measurements section above and the lower phase was carefully transferred to 4 ml liquid scintillation fluid in a scintillation vial and ^14^C activity was measured.

### Liver Histology

Livers were fixed in 10% zinc formalin overnight and then transferred to 70% ethanol for prolonged storage. Tissue processing, sectioning, and staining with hematoxylin and eosin was performed by the Dana Farber/Harvard Cancer Center rodent histopathology core.

### Transmission electron microscopy (TEM)

<u>Whole liver tissue</u>: 8-10 weeks old lean (Wt) and obese (*ob/ob)* mice were anesthetized with 2mg/ml xylazine and 2mg/ml ketamine and perfused with 10mL of saline followed by 10mL of fixative buffer containing 2.5% glutaraldehyde, 2.5% paraformaldehyde in 0.1 M sodium cacodylate buffer (pH 7.4) (Electron Microscopy Sciences, catalog no: 15949). After perfusion, small pieces (1-2 mm^3^) of liver were immersed in the same fixative buffer described above and then sliced at 300-micron thickness with a compresstome (Precisionary Instruments, catalog no: VF-300-0Z). The first 3 slices (~900um) were discarded to reach the full-mid-section cut. The slices were transferred into a fresh fixative solution containing 4 parts of FP stock (2.5 % PFA, 0.06 % picric acid in 0.2M Sodium Cacodylate buffer pH 7.4) and 1 part of 25 % glutaraldehyde and incubated at 4C overnight. Primary hepatocytes: Cells were fixed in a 1:1 ratio in a fixative buffer and Williams Medium. The fixative buffer contained: 4 parts of FP stock (2.5 % PFA, 0.06 % picric acid in 0.2M Sodium Cacodylate buffer pH 7.4) and 1 part of 25 % glutaraldehyde for at least 2h. The tissue slices or cell pellets were then washed in 0.1M cacodylate buffer and post-fixed with 1% Osmiumtetroxide (OsO_4_) / 1.5% Potassiumferrocyanide (K_2_FeCN_6_) for 1 hour, washed two times in water and one time in 50mM maleate buffer pH 5.15 (MB), incubated in 1% uranyl acetate in MB for 1hr followed by 3 washes in MB and subsequent dehydration in grades of alcohol (10min each; 50%, 70%, 90%, 2×10min 100%) followed by incubation for 1hr in propylene oxide. The samples were incubated overnight at 4C in a 1:1 mixture of propylene oxide and TAAB Epon (TAAB Laboratories Equipment Ltd, https://taab.co.uk). The following day, samples were embedded in TAAB Epon and polymerized at 65C for 48 hrs. <u>Sectioning and imaging</u>: ultrathin sections (about 90nm) were generated using a Reichert Ultracut-S microtome and imaged with a JEOL 1200EX electron microscope at 80kV. Images were recorded with an AMT 2K CCD camera. For the whole liver, sections were imaged in low magnification and portal triad and central vein were located. The high magnification images were collected in the intermediary zone, in between the portal and central vein.

### TEM analysis (Parallel and non-parallel organization of ER)

Whole hepatocyte mid-cross-sectional areas were captured at 1500x magnification with a JEOL 80Kb electron microscope. ER was manually annotated as a single pixel wide filament and binary masks of ER were generated with Fiji. Each filament of ER was searched for a nearby neighbor ER filament at the direction perpendicular to the tangent to the curve in a range of 5-20 pixels (corresponds to 55-225 nm distance at 1500x magnification) in Matlab. If more than 50% of the two neighboring ER traces were within the range search, they were accepted as parallel organized. Parallel organized and non-parallel organized ER traces were exported as Tiff images and ratios were calculated from these images using Fiji. The code can be downloaded at https://github.com/gparlakgul

### FIB-SEM sample preparation

300um thick tissue samples were fixed and prepared as described in TEM section. Samples were washed in ice-cold 0.15 M Na-cacodylate buffer for 5 min, 3 times and then incubated in 0.15 M Na-cacodylate solution containing 1% OsO_4_ and 1.5% potassium ferrocyanide for 1h at 4C. Samples were rinsed with water 3 times and incubated for 20 min in 1% thiocarbohydrazide and rinsed again 3 × 5 min with water. Samples were incubated in 2% OsO_4_ for 30 min and then rinsed 3 × 5 min with water, followed by washing with 50mM maleate buffer pH 5.15 (MB) 3 times and incubated overnight at 4C in 1% uranyl acetate in MB. Next day, samples were washed and subsequently dehydrated in grades of alcohol (10min each; 50%, 70%, 90%, 2×10min 100%). Samples were embedded in increasing concentrations of Durcupan resin mixed with ethanol (30min each; 50%, 70%, 90% and 100% Durcupan) followed by a 4 hr incubation in 100% Durcupan. The samples were moved to fresh 100% Durcupan and polymerized at 65C for 24 hrs. Samples were each first mounted to the top of a 1 mm copper post which was in contact with the metal-stained sample for better charge dissipation, as previously described^3^. A small vertical sample post was trimmed to the Region of Interest (ROI) with a width of 100~130 μm and a depth of 90~100 μm in the direction of the ion beam for each sample. The trimming was guided by X-ray tomography data obtained by a Zeiss Versa XRM-510 and optical inspection under a microtome. A thin layer of conductive material of 10-nm gold followed by 100-nm carbon was coated on the trimmed samples using a Gatan 682 High-Resolution Ion Beam Coater. The coating parameters were 6 keV, 200 nA on both argon gas plasma sources, 10 rpm sample rotation with 45-degree tilt.

### 3D large volume FIB-SEM imaging

Samples were imaged sequentially by three customized FIB-SEM systems, each with a Zeiss Capella FIB column mounted at 90° onto a Zeiss Merlin SEM. Details of the Enhanced FIB-SEM systems were previously described^3–5^. Each block face was imaged by a 3nA electron beam with 1.2 keV landing energy at 2 or 3 MHz. A faster SEM scanning rate was applied on samples with stronger staining contrast. The x-y pixel size was set at 8 nm. A subsequently applied focused Ga+ beam of 15nA at 30 keV strafed across the top surface and ablated away 8 nm of the surface. The newly exposed surface was then imaged again. The ablation – imaging cycle continued about once every 60 (3 MHz) to 90 (2 MHz) seconds for two weeks to complete FIB-SEM imaging of one sample. The sequence of acquired images formed a raw imaged volume, followed by post processing of image registration and alignment using a Scale Invariant Feature Transform (SIFT) based algorithm in IMOD, FiJi and MATLAB.

### FIB-SEM segmentation

The organelles, cells and nuclei have been segmented using the 3dEMtrace platform of ariadne.ai ag (https://ariadne.ai/). In brief, ground truth data was generated from the raw FIB-SEM images by manually annotating cells and organelles. Deep convolutional neural networks (CNNs) have been trained using the manually generated ground truth and fine-tuned until a good quality of segmentation was reached for automated segmentation in 3D. The accuracy of the segmentation has been validated by expert inspection. In addition, the accuracy has been quantified by calculating the Jaccard Index or evaluating the object-level recall/precision and f1-score in validation sub-volumes that have not been used for CNN training. Binary Tiff masks were generated for each organelle class separately. For the cells an instance segmentation was provided in order to assign a unique identifier to each cell. Post-segmentation proofreading and correction was done using the open-source software Knossos (https://knossos.app/) and the commercially available Arivis Vision 4D software.

### FIB-SEM ER sheet and tubule sub-segmentation

The segmentation of the sub-compartments of endoplasmic reticulum (ER sheets and ER tubules) was done by using the binary ER masks: 12 images from different layers of each dataset were chosen randomly as training data. ER sheets and ER tubules were annotated manually as separate classes by labeling both lipid bilayer and also filling in the ER luminal space. U-Net-architecture-based machine learning approach has been utilized to train and segment the data by segmenting, for each dataset, each plane using that plane plus the previous and subsequent planes as input. The code can be downloaded at https://github.com/gparlakgul

### FIB-SEM data analysis and visualization

Raw FIB-SEM data and binary Tiff stacks for each organelle class were uploaded to Arivis Vision 4D software as separate channels and ,sis files were generated. Each individual cell and organelle were re-segmented in Vision 4D to create objects. Quantitative measures such as volume, voxel number, surface area and sphericity were calculated. Unspecific objects with less than 10 voxels were eliminated for organelle volume quantification. Cytosolic volumes were calculated by subtracting the the lipid droplet, mitochondria and ER volumes from the total cell volumes. Movies were generated with Arivis Vision 4D, except Fig3C and Fig3D movies, which were generated in Houdini (SideFX) by Ho Man Leung (Refik Anadol Studio, CA).

### Cell culture

Hepa 1-6 cells were cultured in DMEM with 10% CCS. Cos-7 cells were cultured in DMEM with 10% FBS. All cells were cultured at 37°C in a humidified incubator that was maintained at a CO_2_ level of 10%. For the overexpression, Climp-63-Myc plasmid was acquired from Origene (catalog no: MR215622). The plasmid was transfected into Hepa 1-6 and Cos-7 cells using lipofectamine LTX overnight in OptiMEM media and the experiments were done after 36-48h after transfection.

### Primary hepatocyte isolation

Animals were anesthetized using 2 mg/ml xylazine and 2 mg/ml ketamine in PBS. The livers were perfused with 50 mL of buffer I (11 mM Glucose; 200 μM EGTA; 1.17 mM MgSO_4_ heptahydrated; 1.19 mM KH_2_PO_4_; 118 mM NaCl ; 4.7 mM KCl; 25 mM NaHCO_3_, pH 7.32) through the portal vein with an osmotic pump set to the speed of ~4 mL/min until the liver turned pale. The speed was gradually increased until ~7 mL/min afterwards. When the entire buffer I had been infused, it was substituted for 50 mL of buffer II (11 mM Glucose; 2.55 mM CaCl_2_; 1.17 mM MgSO_4_ heptahydrated; 1.19 mM KH_2_PO_4_; 118 mM NaCl ; 4.7 mM KCl; 25 mM NaHCO_3_; BSA (fatty acid free) 7.2 mg/mL; 0.18 mg/mL of Type IV Collagenase (Worthington Biochem Catalog: LS004188), BSA and collagenase were added immediately before use. The buffers were kept at ~37°C during the entire procedure. After perfusion, the primary hepatocytes were carefully released and sedimented at 500 rpm for 2 minutes, washed two times and suspended with Williams E medium supplemented with 5 % CCS and 1 mM glutamine (Invitrogen, CA). To separate live from dead cells, the solution of hepatocytes was layered on a 30% Percoll gradient and centrifuged ~1500 rpm for 15 minutes. The healthy cells were recovered at the bottom of the tube and plated for experimentation.

### Total protein extraction and Immunoblotting

Liver tissues were homogenized in a polytron in cold lysis buffer containing 50 mM Tris-HCl (pH 7.4), 2 mM EGTA, 5 mM EDTA, 30 mM NaF, 10 mM Na_3_VO_4_, 10 mM Na_4_P_2_O_7_, 40 mM glycerophosphate, 1 % NP-40, and 1% protease inhibitor cocktail. After 20-30 minutes incubation on ice, the homogenates were centrifuged at 9000 rpm for 15 minutes to pellet cell debris. The supernatant was removed, and protein concentrations were determined by BCA. The samples were then diluted in 6x Laemmli buffer and heated at 95°C for 5 minutes. The protein lysates were subjected to SDS-polyacrylamide gel electrophoresis, as previously described (Fu et al, 2012). All the immunoblots were incubated with primary antibody overnight at 4°C, followed by incubation with secondary antibody conjugated with horseradish peroxidase (Cell Signaling Technologies) for 1-3 hour at room temperature. Individual membranes were visualized using the enhanced chemiluminescence system (Roche Diagnostics).

### Primary antibody list

<u>anti-</u> Sec61β: Cell Signaling Technology, catalog no: 14648. <u>anti-Sec61α:</u> Cell Signaling Technology, catalog no: 14868. <u>anti-Trapα</u>: Santa Cruz Biotech., catalog no: sc-134987. <u>anti-Calnexin</u>: Santa Cruz Biotech., catalog no: sc-6465. <u>anti-β-tubulin-HRP</u>: Abcam, catalog no: ab21058 <u>anti-Climp-63 (CKAP4)</u>: Novus Biologicals, catalog no: NBP1-26642 and Abcam, catalog no: ab84712. <u>anti-p180 (RRBP1)</u>: Abcam, catalog no: ab95983. <u>anti-Atlastin-3</u>: home-made antibody, generously provided by Dr. Craig Blackstone’s lab at NIH. <u>anti-Reticulon4 (Nogo)</u>: Novus Biologicals, catalog no: NB100-56681. anti-Reep5 (Dp1): Proteintech, catalog no: 14643-1-AP. <u>anti-Ribophorin1</u>: Santa Cruz Biotech., catalog no: sc-12164. <u>anti-Ribophorin2</u>: Santa Cruz Biotech., catalog no: sc-166421. <u>anti-L7a</u>: Cell Signaling Technologies, catalog no: 2415. <u>anti-ATF4</u>: Cell Signaling Technologies, catalog no: 11815. <u>anti-CHOP</u>: Cell Signaling Technologies, catalog no: 2895. <u>anti-PDI</u>: Cell Signaling Technologies, catalog no: 3501. <u>anti-Myc-Tag</u>: Cell Signaling Technologies, catalog no: 2278. <u>anti-Myc-Tag - Alexa Fluor 647</u>: Cell Signaling Technologies, catalog no: 2233. <u>anti-KDEL - Alexa Fluor 488</u>: Abcam, catalog no: ab184819.

### Secondary antibody list

<u>anti-rabbit IgG - HRP</u>: Cell Signaling Technologies, catalog no: 7074. <u>anti-mouse IgG - HRP</u>: Cell Signaling Technologies, catalog no: 7076. <u>anti-goat IgG -HRP</u>: Santa Cruz Biotech, catalog no: sc-2020. <u>anti-rabbit IgG Alexa Fluor 594</u>: Cell Signaling Technologies, catalog no: 8889. <u>anti-rabbit IgG Alexa Fluor 488</u>: Cell Signaling Technologies, catalog no: 4412. <u>anti-rabbit IgG Alexa Fluor 647</u>: Cell Signaling Technologies, catalog no: 4414.

### Gene expression analysis

Tissues were disrupted in Trizol (Invitrogen) using TissueLyser (Qiagen). Chloroform was added to the Trizol homogenates, vortexed thoroughly and centrifuged 12000g for 20 min at 4°C. The top layer was transferred to another tube and mixed with isopropanol and centrifuged again at 12000g for 20 min at 4°C. The RNA found in the precipitate was washed twice with 70% Ethanol and diluted in RNAse free water. Complementary DNA was synthesized using iScript RT Supermix kit (Biorad). Quantitative real-time PCR reactions were performed in triplicates on a ViiA7 system (Applied Biosystems) using SYBR green and custom primers or primer sets based on Harvard Primer Bank. Gene of interest cycle thresholds (Cts) were normalized to TBP housekeeper levels by the ΔΔCt. Primers used are listed in **Supplementary Table 2**.

### Endogenous protein staining and confocal imaging

Primary hepatocytes, Hepa1-6 or Cos-7 cells were seeded on 35 mm round glass bottom imaging dishes in Williams Medium in the presence of 5% CCS (for primary hepatocytes) and DMEM in the presence 10% CCS/FBS overnight at 37°C, 5% CO_2_. The following morning, cells were washed and fixed with 4% paraformaldehyde for 10 min at room temperature (RT) and washed 3x in PBS, before a 20 min permeabilization using 0.2% Triton-×100 in PBS at RT. Primary antibodies were diluted 1:200 for Sec61β antibody, Cell Signaling Technologies (14648) in PBS and the cells were incubated in this solution overnight at 4°C. Next day, cells were washed 3x with PBS, including one long wash for more than 10 min. Secondary antibody was diluted 1:1000 in PBS, and the cells were incubated with it at RT for 1h in the dark. The cells were washed 3x, including one long wash, and if needed, Hoechst was used as nuclear marker, diluted 1:5000 in PBS and incubated for 10 min at RT. Cells were imaged with a Yokogawa CSU-X1 spinning disk confocal system (Andor Technology, South Windsor, CT) with a Nikon Ti-E inverted microscope (Nikon Instruments, Melville, NY), using a 60x or 100x Plan Apo objective lens with a Zyla cMOS camera and NIS elements software was used for acquisition parameters, shutters, filter positions and focus control. Image analysis was performed using Fiji software. Raw single plane Tiff images were analyzed using a Macro and mean intensity values were calculated per image. Fiji Macro can be downloaded at https://github.com/gparlakgul

### Glucose production assay in primary hepatocytes

Isolated primary hepatocytes were plated in 24 well collagen coated plates. Three hours after plating, cells were washed with warm PBS twice and incubated in 0.1% CCS containing Media 199 (Thermo Scientific cat: 11150067) and with the corresponding adenovirus (LacZ or Climp-63-myc at 30 MOI dose) overnight. Next morning, cells were washed with PBS and incubated in DMEM with no glucose, no glutamine (Gibco catalog no: A1443001) for 1 hour. Then, cells were treated with gluconeogenic substrates in DMEM (A1443001) at a final concentration of: Lactate: 4.5mM, Na-pyruvate: 0.5mM, Glutamine: 2.5mM, or with Glycerol: 20mM, all in the presence of Glucagon (100nM). 3 hours later supernatant was collected, and glucose levels were measured using the Amplex Red Glucose Oxidase Assay Kit (A22189). Plate was washed with PBS and frozen for protein measurement.

### ASGR assay

The ASGR reporter plasmid described in^6^ was cloned in an Adenovirus Ad5 backbone from Vector BioLabs. The adenovirus (serotype 5, Ad5) was generated and amplified by Vector Biolabs. For the experiments in primary hepatocytes, cells were infected with 30 MOI in 12 or 24 well plates with Ad-ASGR and Ad-LacZ or Ad-Climp-63 viruses. The next day, cells were changed to a fresh medium containing phenol-red-free Williams E supplemented with 5 % CCS and 1 mM glutamine (Invitrogen, CA) and incubated for 22 to 24 hours. 10 ul of supernatant was transferred to two 96-well white plates for luciferase assays following the manufacturer’s protocol. Briefly, 50 ul of luciferase substrate (1 uM Cypridina or 10 mM CTZ in 100 mM tris buffer, pH 7.5) was added to the 10 ul medium and incubated in the dark for 5 to 10 min. The luminescence was read on SpectraMax Paradigm plate reader (Molecular Devices).

### Total ER and rough and smooth ER isolation

<u>For the total microsome isolation:</u> subcellular fractionation of the liver was done based on published protocols^1,7^. Briefly, mice were sacrificed and 1g of liver was immediately weighed and washed in cold Buffer 1 (content described below). The tissue was minced and immersed in Buffer2. The tissue was filtered in order to eliminate the blood and transferred to a glass potter in 15mL of Buffer1 and further disrupted by Dounce homogenization. The homogenate was spun down at 740*g* for 5 min twice in a low-speed centrifuge; the supernatant was recovered and further centrifuged for 10 min at 10,000*g,* 3 times. The resulting pellet (crude mitochondrial fraction) was discarded, and the supernatant was used for the collection of the ER fraction which was obtained by centrifuging the supernatant at 100.000g for 60 min. <u>Buffer1</u>: Mannitol 225mM, Sucrose 75mM, Tris-HCl 30mM, BSA 0.5% and EGTA 0.5mM, pH7.4. <u>Buffer2</u>: Mannitol 225mM, Sucrose 75mM, Tris-HCl 30mM, BSA 0.5%, pH 7.4. Buffer3: Mannitol 225mM, Sucrose 75mM, Tris-HCl 30mM, pH 7.4. <u>For the rough and smooth ER fractionation</u>: the fractionation protocol was based on^8^ with modifications. Briefly, mice were sacrificed, the liver was excised, weighted and washed in a petri dish containing ice-cold sucrose buffer. Liver was transferred to a beaker containing ice-cold fresh sucrose buffer and minced in small pieces. The buffer was removed through a mesh filter and a volume of sucrose buffer was added an equivalent to ~5 times volume to the weight of the livers. The liver pieces were then transferred to a 15mL Dounce homogenizer with a Teflon pestle and homogenized with 5 strokes. The mixture was centrifuged 2 times at 10.000g for 20 min in a slow speed centrifuge, rotor: JA-17. The pellet containing mitochondria, and other contaminants were discarded. CsCl was added to the supernatant to make a final concentration of 15mM CsCl with using 1M CsCl stock solution. The supernatant was then layered on top of 15ml of 1.3M sucrose + 15mM CsCl in an Beckman SW32 Ti centrifuge tube (Cat No: 344058).The solution was completed to 34ml.s with 0.25M sucrose buffer and centrifuged for 4 hours at 105,000g, 4°C (with Beckman SW32 rotor). The pellet will contain rough microsomes and the interface between 0.25-1.3M will contain smooth microsomes. The supernatant above the band was drawn off (soluble fraction) and saved on ice and the pellet was resuspended in 0.25 M sucrose by gentle homogenization in a glass homogenizer. Both subfractions were centrifuged in a Ti50 rotor at 225,000g for 1h (Beckman Type 70.1Ti rotor) at 4C. Final pellets were resuspended in 0.25 M sucrose. <u>Sucrose buffer</u>: 250mM Sucrose and 50mM Tris pH7.4 + protease inhibitor cocktail, pH adjusted to 7.2 at 4°C.

### Statistical Analysis

Statistical significance was assessed using GraphPad Prism Version 7, using the Student’s t-test and P values are indicated in the figure legends. Two-way ANOVA was used to evaluate significance in **Fig. 4d, f**. All data are mean ± SEM.

## Acknowledgements

We are especially grateful for Ho Man Leung, Nicholas Boss and Refik Anadol for their artistic talent and vision, visualization of the data, generating videos and generously sharing the resources and expertise of the Refik Anadol Studio (Los Angeles, CA). We would like to thank Elizabeth Benecchi, Maria Ericsson and Louise Trakimas for their help in sample preparation for TEM. We appreciate Marcelo Cicconet for his help with generating the Matlab codes. We would like to thank Christopher Zugates for his help and guidance in using the Arivis Vision 4D software. We appreciate Adrian Wanner, Jörgen Kornfeld and the whole Ariadne team’s effort and help with the segmentation. We thank all members of the Sabri Ülker Center and Hotamisligil Lab community for their continued support and encouragement.

## Author contributions

G.P. and A.P.A. formulated the questions, designed the project and performed the *in vitro* and *in vivo* experiments, analyzed the data, prepared the figures, and wrote and revised the manuscript. E.C., E.G, N.M, Y.L performed and assisted in vitro and in vivo experiments and part of the imaging analysis. S.P., C.S.X, and H.F.H. performed, supervised and executed collection of the FIB-SEM data. G.S.H. conceived, supervised and supported the project, designed experiments, interpreted results, and revised the manuscript.

## Data availability

Supplementary videos can be reached at https://www.youtube.com/playlist?list=PLpzquMkvsJ9XZvqAKyHcz62iDQkoKxmpx

## Code availability

The source code generated and analyzed during this study can be found at https://github.com/gparlakgul

## Additional information

Supplementary Information file includes methods, Supplementary videos 1-9 and Supplementary Tables 1–2

## REFERENCES

1. Ben-Moshe, S. & Itzkovitz, S. Spatial heterogeneity in the mammalian liver. Nat. Rev. Gastroenterol. Hepatol. 16, 395–410 (2019).

2. Lee, A. H., Chu, G. C., Iwakoshi, N. N. & Glimcher, L. H. XBP-1 is required for biogenesis of cellular secretory machinery of exocrine glands. EMBO J. 24, 4368–4380 (2005).

3. Zirkin, B. R. & Papadopoulos, V. Leydig cells: Formation, function, and regulation. Biology of Reproduction vol. 99 101–111 (2018).

4. Valm, A. M. et al. Applying systems-level spectral imaging and analysis to reveal the organelle interactome. Nature vol. 546 162–167 (2017).

5. Lee, J. E., Cathey, P. I., Wu, H., Parker, R. & Voeltz, G. K. Endoplasmic reticulum contact sites regulate the dynamics of membraneless organelles. Science (80-.). 367, (2020).

6. West, M., Zurek, N., Hoenger, A. & Voeltz, G. K. A 3D analysis of yeast ER structure reveals how ER domains are organized by membrane curvature. J. Cell Biol. 193, 333–346 (2011).

7. Nixon-Abell, J. et al. Increased spatiotemporal resolution reveals highly dynamic dense tubular matrices in the peripheral ER. Science (80-.). 354, (2016).

8. Heinrich, L. et al. Automatic whole cell organelle segmentation in volumetric electron microscopy. bioRxiv 2020.11.14.382143 (2020) doi:10.1101/2020.11.14.382143.

9. Xu, C. S. et al. Enhanced FIB-SEM systems for large-volume 3D imaging. Elife 6, (2017).

10. Xu, C.S., Hayworth K.J., H. H. F. Enhanced FIB-SEM systems for large-volume 3D imaging. (2020). US Patent 10,600,615.

11. Xu, C. S., Pang, S., Hayworth, K. J. & Hess, H. F. Transforming FIB-SEM Systems for Large-Volume Connectomics and Cell Biology. in Neuromethods vol. 155 221–243 (Humana Press Inc., 2020).

12. Goyal, U. & Blackstone, C. Untangling the web: Mechanisms underlying ER network formation. Biochim. Biophys. Acta - Mol. Cell Res. 1833, 2492–2498 (2013).

13. Shibata, Y. et al. Mechanisms determining the morphology of the peripheral ER. Cell 143, 774–788 (2010).

14. Chen, S., Novick, P. & Ferro-Novick, S. ER structure and function. Current Opinion in Cell Biology vol. 25 428–433 (2013).

15. Westrate, L. M., Lee, J. E., Prinz, W. A. & Voeltz, G. K. Form follows function: The importance of endoplasmic reticulum shape. Annual Review of Biochemistry vol. 84 791–811 (2015).

16. Terasaki, M. et al. Stacked Endoplasmic Reticulum Sheets Are Connected by Helicoidal Membrane Motifs. Cell 154, 285–296 (2013).

17. Lynes, E. M. & Simmen, T. Urban planning of the endoplasmic reticulum (ER): How diverse mechanisms segregate the many functions of the ER. Biochimica et Biophysica Acta - Molecular Cell Research vol. 1813 1893–1905 (2011).

18. Voeltz, G. K., Prinz, W. A., Shibata, Y., Rist, J. M. & Rapoport, T. A. A class of membrane proteins shaping the tubular endoplasmic reticulum. Cell 124, 573–586 (2006).

19. Sandoz, P. A. & Van Der Goot, F. G. How many lives does CLIMP-63 have? Biochemical Society Transactions vol. 43 222–228 (2015).

20. Shen, B. et al. Calumenin-1 Interacts with Climp63 to Cooperatively Determine the Luminal Width and Distribution of Endoplasmic Reticulum Sheets. iScience 22, 70–80 (2019).

21. Zhang, H. & Hu, J. Shaping the Endoplasmic Reticulum into a Social Network. Trends in Cell Biology vol. 26 934–943 (2016).

22. Fu, S. et al. Phenotypic assays identify azoramide as a small-molecule modulator of the unfolded protein response with antidiabetic activity. Sci. Transl. Med. 7, (2015).

23. Klopfenstein, D. R. et al. Subdomain-specific localization of CLIMP-63 (p63) in the endoplasmic reticulum is mediated by its luminal α-helical segment. J. Cell Biol. 153, 1287–1299 (2001).

24. Vedrenne, C., Klopfenstein, D. R. & Hauri, H. P. Phosphorylation controls CLIMP-63-mediated anchoring of the endoplasmic reticulum to microtubules. Mol. Biol. Cell 16, 1928–1937 (2005).

25. Terasaki, M., Chen, L. B. & Fujiwara, K. Microtubules and the endoplasmic reticulum are highly interdependent structures. J. Cell Biol. 103, 1557–1568 (1986).

26. Noda, C. et al. Valosin-containing protein-interacting membrane protein (VIMP) links the endoplasmic reticulum with microtubules in concert with cytoskeleton-linking membrane protein (CLIMP)-63. J. Biol. Chem. 289, 24304–24313 (2014).

27. Nikonov, A. V., Hauri, H. P., Lauring, B. & Kreibich, G. Climp-63-mediated binding of microtubules to the ER affects the lateral mobility of translocon complexes. J. Cell Sci. 120, 2248–2258 (2007).

28. Shemesh, T. et al. A model for the generation and interconversion of ER morphologies. Proc. Natl. Acad. Sci. 111, E5243–E5251 (2014).

29. Zhu, Y. et al. Sec61β facilitates the maintenance of endoplasmic reticulum homeostasis by associating microtubules. Protein Cell 9, 616–628 (2018).

## METHOD REFERENCES

1. Fu, S. et al. Aberrant lipid metabolism disrupts calcium homeostasis causing liver endoplasmic reticulum stress in obesity. Nature 473, 528–531 (2011).

2. Arruda, A. P. et al. Defective STIM-mediated store operated Ca2+ entry in hepatocytes leads to metabolic dysfunction in obesity. Elife 6, (2017).

3. Xu, C. S. et al. Enhanced FIB-SEM systems for large-volume 3D imaging. Elife 6, (2017).

4. Xu, C.S., Hayworth K.J., H. H. F. Enhanced FIB-SEM systems for large-volume 3D imaging. (2020). US Patent 10,600,615.

5. Xu, C. S., Pang, S., Hayworth, K. J. & Hess, H. F. Transforming FIB-SEM Systems for Large-Volume Connectomics and Cell Biology. in Neuromethods vol. 155 221–243 (Humana Press Inc., 2020).

6. Fu, S. et al. Phenotypic assays identify azoramide as a small-molecule modulator of the unfolded protein response with antidiabetic activity. Sci. Transl. Med. 7, (2015).

7. Arruda, A. P. et al. Chronic enrichment of hepatic endoplasmic reticulum-mitochondria contact leads to mitochondrial dysfunction in obesity. Nat. Med. 20, 1427–1435 (2014).

8. Margolis, R. N., Cardell, R. R. & Curnow, R. T. Association of glycogen synthase phosphatase and phosphorylase phosphatase activities with membranes of hepatic smooth endoplasmic reticulum. J. Cell Biol. 83, 348–356 (1979).

